# Molecular Insights into CLD Domain Dynamics and Toxin Recruitment of the HlyA *E. coli* T1SS

**DOI:** 10.1101/2025.05.27.656419

**Authors:** Rocco Gentile, Stephan Schott-Verdugo, Sakshi Khosa, Michele Bonus, Jens Reiners, Sander H. J. Smits, Lutz Schmitt, Holger Gohlke

**Author notes:** Prof. Dr. Holger Gohlke: Institute for Pharmaceutical and Medicinal Chemistry, Heinrich Heine University Düsseldorf, Universitätstr. 1. 40225 Düsseldorf, Germany, and Institute of Bio– and Geosciences (IBG-4: Bioinformatics), Forschungszentrum Jülich GmbH, Wilhelm-Johnen-Str., 52425 Jülich, Germany, **Email:** or. **Author Contributions** R.G.: conceptualization, computational investigation, analysis, and visualization. S.S.-V. and M.B.: computational investigation, supervision, analysis. S.K. and J.R.: experimental investigation, analysis, and visualization; S.H.J.S., L.S., and H.G.: conceptualization, supervision, analysis, funding, resources, and project management. The manuscript was written with contributions of all authors. All authors have given approval for the final version of the manuscript. **Competing Interest Statement** The authors declare no competing financial interest.

## Abstract

*Escherichia coli* is a Gram-negative opportunistic pathogen causing nosocomial infections through the production of various virulence factors. Type 1 secretion systems (T1SS) contribute to virulence by mediating one-step secretion of unfolded substrates directly into the extracellular space, bypassing the periplasm. A well-studied example is the hemolysin A (HlyA) system, which secretes the HlyA toxin in an unfolded state across the inner and outer membranes. T1SS typically comprise a homodimeric ABC transporter (HlyB), a membrane fusion protein (HlyD), and the outer membrane protein TolC. Some ABC transporters in T1SS also contain N-terminal C39 peptidase or peptidase-like (CLD) domains implicated in substrate interaction. Recent cryo-EM studies have resolved the inner-membrane complex as a trimer of HlyB homodimers with associated HlyD protomers. However, a full structural model including TolC remains unavailable. We present the first complete structural model of the HlyA T1SS, constructed using template– and MSA-based information and validated by SAXS. Molecular dynamics simulations provide insights into the function of the CLD domains, which are partially absent from existing cryo-EM structures. These domains may modulate transport by stabilizing specific conformations of the complex. Simulations with a C-terminal fragment of HlyA indicate that toxin binding occurs in the occluded conformation of HlyB, potentially initiating substrate transport through a single HlyB protomer before transitioning to an inward-facing state. HlyA binding also induces allosteric effects on HlyD, altering key residues involved in TolC recruitment. These results indicate how substrate recognition and transport are coupled and may support the development of antimicrobial strategies targeting the T1SS.

## Introduction

Type 1 secretion systems (T1SS) are highly efficient protein translocation systems in Gram-negative bacteria, capable of secreting substrates ranging from 20 to 1,500 kDa (1–5). Due to the variability in substrate length, the conventional alternating access transport mechanism of transport proteins (6) is unlikely to be applicable. Furthermore, the secretion signal is encoded at the extreme C-terminus of the substrate (7, 8). Consequently, secretion initiates only after ribosomal synthesis of the entire polypeptide, with the C-terminus appearing first from the bacterial cell in a vectorial manner (9). Therefore, the mechanism of T1SS-mediated substrate secretion must differ from that of the well-characterized Sec translocon (10).

Among T1SS, the *Escherichia coli* hemolysin A (HlyA) transport system has long served as the model to study T1SS (5, 11). Recent cryo-EM experiments have revealed the structure of the inner membrane complex (IMC) of the HlyA transport system. HlyA is a 110-kDa pore-forming toxin capable of lysing human cells (12). The toxin is secreted in an unfolded state from the cytosol into the extracellular space, where extracellular Ca^2+^ induces its folding (13, 14). Cryo-EM structures show that the IMC consists of three HlyB homodimers and six HlyD subunits forming a heterododecameric assembly (15). HlyB is a member of the ATP-binding cassette (ABC) superfamily of transporters, which are molecular pumps powered by ATP hydrolysis. The relevance of ATP hydrolysis in HlyA secretion is supported by findings that mutations in HlyB that abolish ATP hydrolysis also prevent HlyA secretion (16). Like all ABC transporters, HlyB contains two transmembrane domains (TMDs) and two nucleotide-binding domains (NBDs). HlyD is a periplasmic adaptor protein unique to Gram-negative bacteria. It contains a small N-terminal cytosolic domain, a single transmembrane (TM) helix, and a large periplasmic domain that interacts with TolC (17–19). Proteins homologous to HlyD are also found in tripartite drug efflux systems, where they form a critical link between the inner membrane transporter and the outer membrane porin (20), which is TolC in *E. coli*. TolC forms a homotrimeric complex that spans the outer membrane and part of the periplasm, connecting with the periplasmic tip of HlyD to create a continuous conduct for the export of HlyA (17, 21). Binding of the HlyA substrate to HlyB has been suggested to trigger conformational changes that propagate through HlyD, promoting its engagement with TolC only when HlyA is present (22).

Functional studies indicate that HlyA variants are translocated via a canonical ABC transport pathway after binding to the N-terminal C39 peptidase-like domain (CLD) of HlyB, a region homologous to C39 cysteine proteases but lacking the catalytic cysteine residue and, consequently, being enzymatically inactive (23). The cryo-EM structures reveal that only one of the two CLDs (CLD1) is structured and positioned between neighboring protomers, irrespective of the occupation of the ATP binding site. They also show that neighboring HlyB protomers form contacts via their NBDs, and the six HlyD subunits link adjacent HlyB dimers. In the ATP-bound form, all HlyB protomers exhibit the same conformation (occluded with the NBDs in the ATP-bound dimeric state), while in the nucleotide-free form, one dimer displays an inward-facing, NBD-separated conformation. These findings, together with functional studies, have led to a model of HlyA secretion, in which the substrate is recruited into the translocation pathway of one HlyB, while the other two hydrolyze ATP to power the translocation (15). A central pore is located at the interface of the three dimers, hydrophobic, and partially filled with lipid molecules. The asymmetric structure of HlyB dimers within the trimeric complex raises the question of whether all three HlyB dimers bind to and hydrolyze ATP. To address this, the ATP hydrolysis rate of the HlyB/D complex was compared with that of the isolated HlyB dimer. These data suggest that all three HlyB dimers in the inner membrane complex can bind and hydrolyze ATP with no apparent cooperativity between them (15). However, despite these structural and functional insights, the structure of the tripartite T1SS complex and atomistic insights into toxin binding to HlyB and details underlying subsequent TolC recruitment have remained elusive.

Here, we address the single dimer transport hypothesis, considering that many mechanistic details of substrate translocation still remain unclear. We combined template– and MSA-based co-folding structure prediction techniques to generate a complete heterododecameric HlyB/HlyD complex with differing HlyB states, assess their structural variability with all-atom molecular dynamics (MD) simulations, generate and assess structural models of a fragment of HlyA, HlyA2, binding to HlyB, and scrutinize with a model of dynamic allostery (24–27) how such binding can influence regions in HlyD critical for recruiting TolC and extend into TolC. The modeling results are supported by SAXS data of a stalled tripartite T1SS.

## Results

### MD simulations of HlyB/HlyD complexes reveal the least structural variability of functionally relevant parts in HlyBOOO/HlyD and HlyBIOO/HlyD

Two cryo-EM structures of HlyB/HlyD complexes have been resolved (PDB IDs: 7SGR and 8DCK) (15), one (8DCK) with ATP/Mg^2+^ bound to HlyB. Here, all HlyB dimers adopted an occluded conformation. In the other structure without ATP/Mg^2+^ (7SGR), only one HlyB dimer is in an inward-facing conformation. We refer to these states as HlyB**_OOO_** and HlyB**_IOO_**, respectively. The transmembrane (TM) region of each HlyB protomer is preceded by a cytoplasmic C39 peptidase-like domain (CLD) domain. The solvent-exposed CLD (termed CLD2) has been hypothesized to bind hemolysin but is not resolved in either cryo-EM structure. The other one (termed CLD1) is buried by the protomer neighboring a HlyB dimer in the trimeric assembly of HlyB dimers and ordered. HlyD is also considerably fragmented in the cryo-EM maps. Thus, a complete experimentally resolved structural model of the HlyB/HlyD complex is not yet available.

We complemented the missing parts in the two cryo-EM structures by using Colabfold (28) with these structures as templates, and MODELLER (29, 30) for tethering the models with the available structures and modeling connecting loops. That way, we generated complete heterododecameric HlyB_OOO_/HlyD and HlyB_IOO_/HlyD complexes with all HlyB dimers in the occluded conformation (HlyB_O_) or one of them in the inward-facing conformation (HlyB_I_) (Table S1). To expand the structural repertoire of HlyB/HlyD complexes, we also generated complete models with two or three HlyB dimers in the inward-facing conformation (HlyB_IIO_/HlyD, HlyB_III_/HlyD; Table S1). Each one of the four systems was embedded in a DOPE:DOPG 3:1 membrane bilayer resembling the inner membrane of Gram-negative bacteria, especially *E. coli* (31), and subjected to unbiased molecular dynamics simulations for 1 μs length, using five independent replicas in each case. All unbiased MD simulations showed membrane phases that correspond with those generally found by experiments and MD simulations for membranes of Gram-negative bacteria (32) (Fig. S5).

The MD simulations revealed pronounced differences in the structural variabilities of the complex components (**Figure 1A-E**), and these differences are generally seen across the replicas. In the case of HlyB_III_/HlyD, both HlyB_I_ and HlyD reveal large RMSD values, which are mirrored by the almost complete lack of occupancy densities for CLD1 and CLD2. The occupancy density is the time-averaged probability density of the spatial occupancy of a given protein part, derived from the MD trajectories. In this complex, also part of the NBD structure varies, which leads to large RMSD values of the bound ATP/Mg^2+^ (**Figure S6**). CLD1 of HlyB_O_ also showed high structural variability in the case of HlyB_IIO_/HlyD, contrary to what has been found for the well-resolved domains in HlyB_O_ in the cryo-EM structures. In both HlyB_OOO_/HlyD and HlyB_IOO_/HlyD, the structural variabilities of CLD1 are the lowest, and this is independent of the HlyB conformation in the latter complex. Furthermore, the CLD2 domains within the HlyB_IOO_/HlyD show markedly lower structural variability than those in HlyB_OOO_/HlyD. Notably, within HlyB_IOO_/HlyD, the presence of the single HlyB_I_ appears to stabilize the CLD2 domains within the two HylB_O_ dimers. In contrast, configurations with more than one HlyB_I_ dimer lead to increased positional variability of CLD2. Overall, CLD1 remains the least variable in both HlyB_OOO_/HlyD and HlyB_IOO_/HlyD. The CLD2 domain, functionally relevant for toxin translocation (15, 33), also shows the lowest variability in HlyB_IOO_/HlyD, the arrangement also found in the cryo-EM structures. Consequently, we selected these two configurations (HlyB_OOO_/HlyD and HlyB_IOO_/HlyD) for the subsequent analyses.

**Figure 1.**
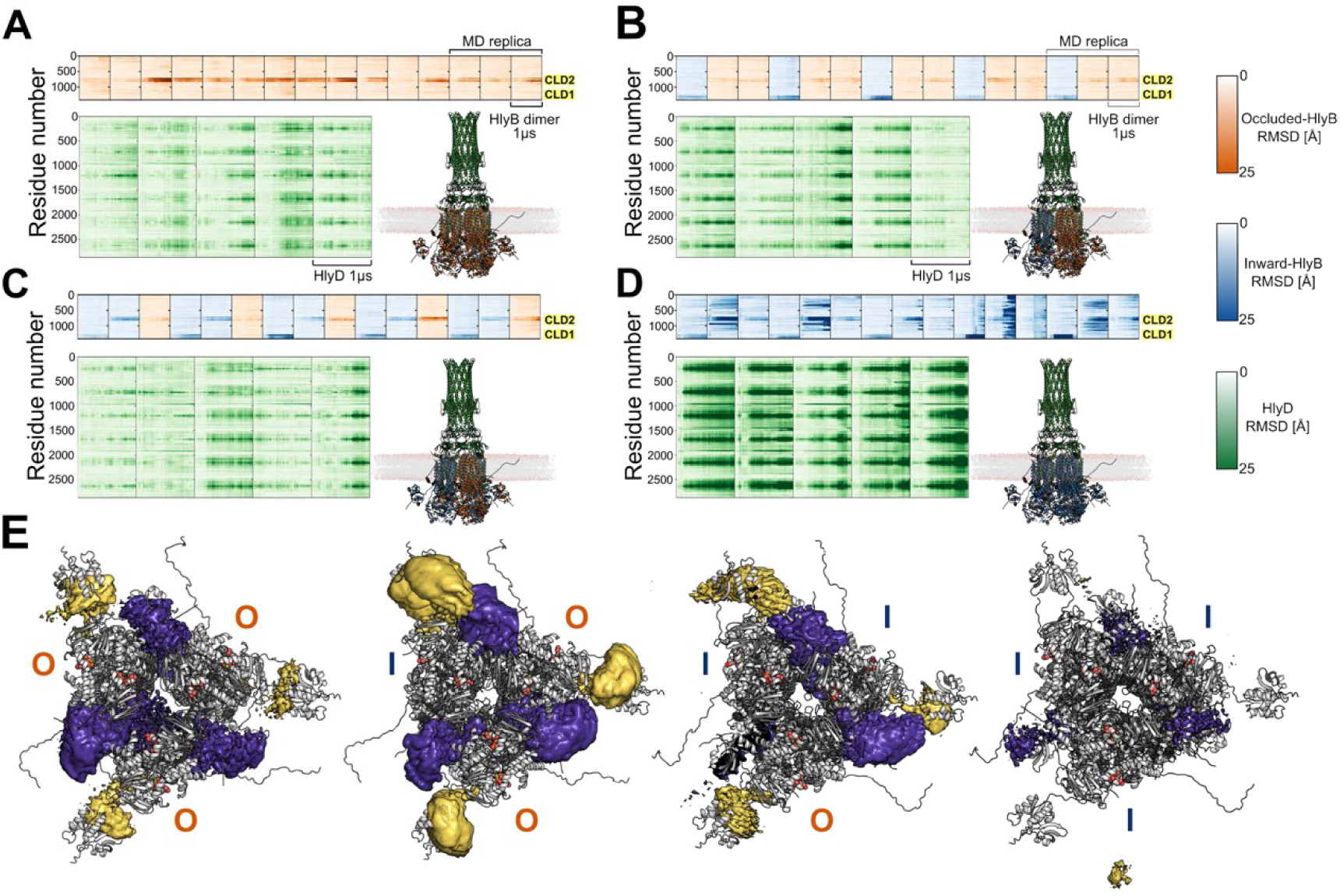
Structural variabilities of motifs in different complex configurations. Best-fit per-residue RMSDs onto the first frame of production runs for **A)** HlyB_OOO_/HlyD, **B)** HlyB_IOO_/HlyD, **C)** HlyB_IIO_/HlyD, **D)** HlyB_III_/HlyD. Each panel depicts data for occluded HlyB units in orange, for inward-facing HlyB units in blue, and for HlyD in green. Structures are colored accordingly. All complexes were inserted in a DOPE:DOPG 3:1 membrane bilayer and simulated in five replicas for 1 μs length. **E)** Average density maps of CLD1 (purple) and CLD2 (yellow) for the four depicted configurations (full-occluded O-O-O, single-inward I-O-O, double inward I-I-O, and full-inward I-I-I) computed with CPPTRAJ (71). All atoms were considered in the 3D density grid calculations, and the contour level was set as 1 standard deviation above the mean value (1σ).

### Structural models of HlyA2 binding, impact on CLD2 structural variability and HlyB interactions

The CLD domain is essential for HlyA secretion, but only one domain per HlyB dimer (CLD2) in the IMC is exposed to the cytoplasm and available to bind the toxin. HlyA binds to the CLD domain with its RTX domain (33) (**Figure 2A**). Lecher et al. (23) used a truncated HlyA fragment, HlyA2 (residues 807-966). HlyA2 is known to interact with the CLD through GG motifs (GGxGxDxUx, where U is a large lipophilic residue and x any aminoacid) of the RTX domain and to fold in the presence of a Ca^2+^ concentration above 500 μM (23), present extracellularly. Intracellularly, where Ca^2+^ concentrations are lower, the toxin is recruited and transported in an unfolded state (34), which poses challenges for template-, ab initio-, or coevolutionary-based structure prediction tools used for complex model generation. When using AlphaFold3 (35, 36) to model the interaction between HlyA and CLD, HlyA2 was predicted to mostly contain β-strands (**Figure S3A, B**). Lecher et al. demonstrated that only the unfolded state of the substrates, HlyA1 or HlyA2, interacted with the isolated CLD while the folded states did not show any interactions (23). Furthermore, the model scores indicated that the model quality is low, particularly the interface predicted template modeling (ipTM) score (**Figure S3C**).

**Figure 2.**
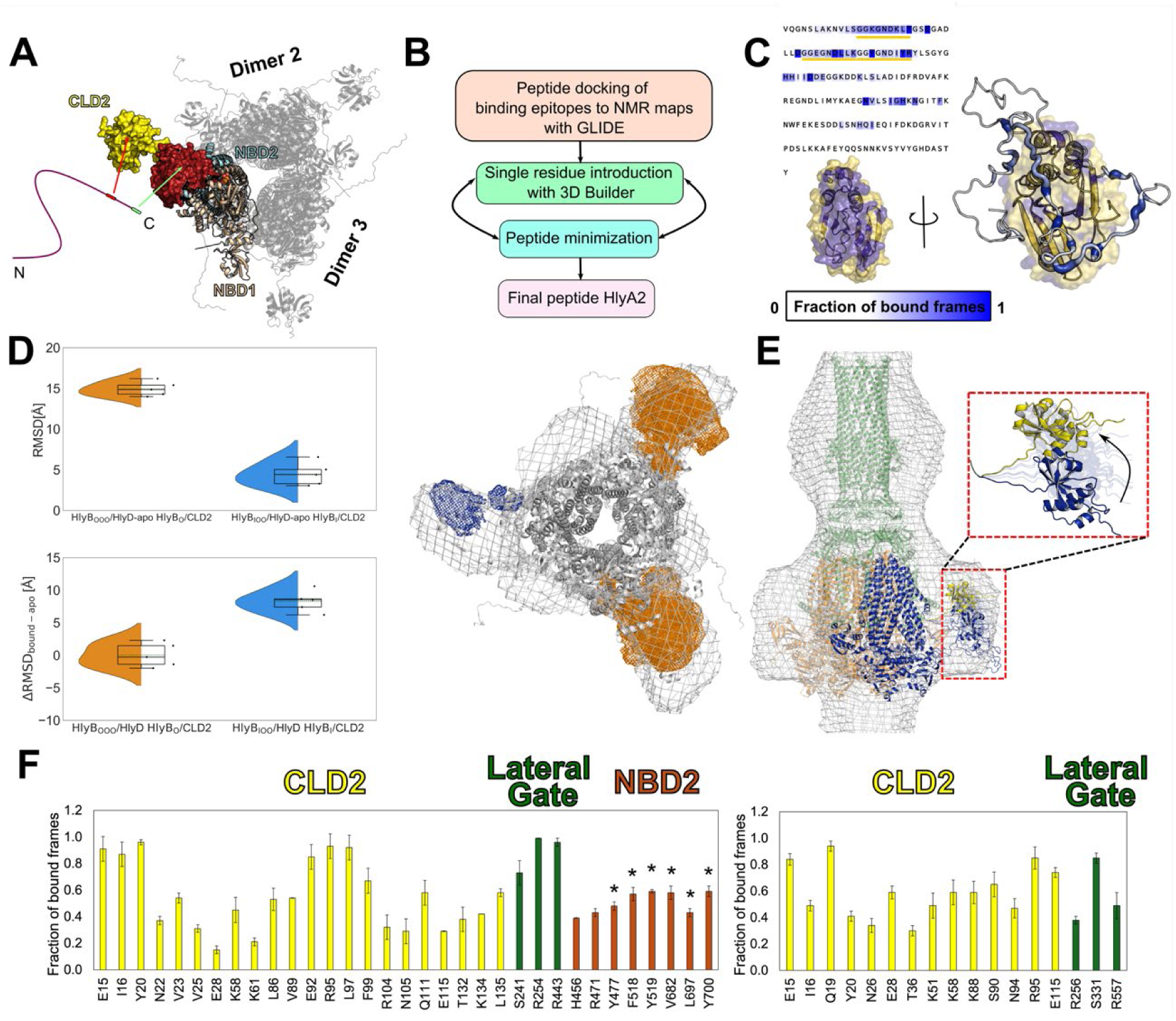
**HlyA2 modeling and binding to CLD2 of HlyB**_OOO_**/HlyD or HlyB**_IOO_**/HlyD. A)** A scheme of the interaction motifs of HlyA2 for binding to HlyB. Adapted from ref. (33). **B)** The workflow explaining how HlyA2 binding to the CLD2 was modeled using Schrödinger software tools (39). Each single-residue introduction was followed by peptide minimization until the full-length HlyA2 (residues 807-966) was obtained. **C)** Chemical shift changes described in ref. (23) upon HlyA binding were mapped onto the CLD2 structure. Interacting regions are highlighted in yellow, while noninteracting regions are shown in purple. A representative HlyA2 conformation is shown as a putty cartoon representation around CLD2. The putty cartoon width and intensity of the blue color indicate a prolonged interaction during MD simulations (see color scale). The interaction persistence was also mapped at the sequence level. The GG motifs show the most prolonged interactions with the CLD2 (underlined in yellow). **D)** RMSD of HlyA2-bound CLD2 with respect to the first frame of each trajectory (left). The violin plots indicate the distribution of the data points; the inner box of a box plot represents the interquartile range, with the horizontal black line indicating the median. The dotted green line represents the mean, and the whiskers show the rest of the distribution, excluding points determined to be outliers when they fall outside 1.5 times the interquartile range. The orange violins indicate a CLD2 in (HlyA2)/HlyB_OOO_/HlyD and the blue violins the CLD2 of HlyB_I_ in (HlyA2)/HlyB_IOO_/HlyD. Top: No HlyA2 was bound. Bottom: HlyA2 was bound; ΔRMSD_bound-apo_ is reported. **E)** *Left:* 3D density grid illustrating the distribution of CLD2 domains computed with CPPTRAJ (71) for the HlyB_IOO_/HlyD from 5 replicas of 1 μs length and their fitting in the SAXS 3D shape reconstruction of the eGFP-HlyA/HlyB/HlyD/TolC complex (grey mesh). All atoms were considered in the 3D density grid calculations, and the contour level was set as 1 standard deviation above the mean value (1σ). CLD2 from HlyB_O_ in HlyB_IOO_/HlyD is colored in orange; CLD2 from HlyB_I_ in HlyB_IOO_/HlyD is colored in blue. The SAXS 3D shape reconstruction is depicted with z-clipping at the level of 150 Å from the top view. Note that a three-fold rotation symmetry was applied for generating the SAXS 3D shape reconstruction, which therefore does not distinguish between CLDs in apo versus bound states. *Right:* SAXS 3D shape reconstruction of the eGFP-HlyA/HlyB/HlyD/TolC complex (grey mesh) viewed from the side, with the HlyA2/HlyB_IOO_/HlyD fitted into it. HlyB_O_ is depicted in orange, HlyB_I_ in blue, and HlyD in green. The red-dashed box shows an overlay of the starting structure (blue cartoon), final averaged structure (yellow cartoon), and selected ensemble members (transparent ribbon) across the five MD simulation trajectories. The movements of HlyA2-bound CLD2 (red-dashed box) fit with the spacious SAXS 3D shape reconstruction of this region. **F)** Interaction persistence expressed as the fraction of frames with HlyB residues interacting with HlyA2. The *left* plot indicates the binding of HlyA2 to the CLD2 of HlyB_OOO_/HlyD and the *right* plot shows the binding to the CLD2 of HlyB_I_ in HlyB_IOO_/HlyD. CLD2 residues are shown as yellow histogram bars, lateral gate residues in green, and NBD2 residues in orange. The standard error of the mean (SEM) with *n* = 5 replicas is depicted. NBD residues predicted to be important for HlyA interaction in ref. (33) are marked with a *.

Therefore, we adopted a stepwise approach for modeling HlyA2 binding to CLD (**Figure 2B**). Known interacting epitopes on the CLD (**Figure 2A, C**) (23) were used as constraints to guide the docking of known interacting HlyA2 peptide fragments (33) to CLD with Glide (37, 38). Missing residues were added one after the other using the 3D Builder tool (39), followed by peptide minimization after each addition, until a model of the HlyA2/CLD complex was obtained. The resulting HlyA2/CLD complex was solvated and simulated without restraints for 1 μs in five replicas. The results showed that the described binding motifs (23) were consistently maintained (**Figure 2C**).

The toxin is known to also interact with the NBD domain of HlyB (33) (**Figure 2A**). To assess if HlyA2 is able to interact with HlyB in other regions besides the CLD, two further models were generated: one HlyA2 bound to one CLD2 of HlyB_OOO_/HlyD and one HlyA2 bound to the CLD2 of HlyB_I_ in HlyB_IOO_/HlyD. Either system was embedded in a DOPE:DOPG 3:1 membrane (31) and solvated as done for the apo-complexes and simulated for 1 μs in five replicas. While the structural variability of the CLD2 in HlyB_O_ remains largely unaffected by the binding of HlyA2, it increases for the CLD2 of HlyB_I_ if HlyA2 is bound (**Figure 2D**).

Fusing HlyA with the fast-folding eGFP results in a stalled T1SS, where eGFP-HlyA, HlyB, and HlyD were overexpressed while endogenous TolC completed the T1SS complex (16, 40). In this study, we used the concept of a stalled T1SS, with a His_6_ tag and eGFP at the N-terminus and FLAGx3-L tag (see Materials and Methods section) at the C-terminus of the RTX domain (before the secretion signal), respectively, of HlyA for purification purposes. The stalled T1SS was purified via the Flag tag and eluted as a homogenous peak in the size exclusion chromatography (F**igure S7**). The presence of all the components of the Hly T1SS (HlyA, HlyB, HlyD, and TolC) could be verified through Western blot analysis. The ab initio 3D shape reconstruction of the stalled eGFP-HlyA/HlyB/HlyD/TolC complex determined by SAXS revealed pronounced density regions that overlap well with occupancy densities of CLD2 domains determined from MD simulations of HlyA2/HlyB_IOO_/HlyD (**Figure 2E**, left). The occupancy density for the HlyA2-bound CLD2 of HlyB_I_ is less distinctive than those for CLD2s of HlyB_O_, in line with the increased structural variability observed in MD simulations for the former case. The SAXS 3D shape reconstruction has a threefold rotation symmetry, as imposed during the process of the SAXS ab initio model generation. This implies that putative differences between toxin-bound and –unbound CLDs in the stalled experimental complex cannot be detected. Still, the SAXS 3D shape reconstruction confirms the observation from the cryo-EM structures that one of the CLDs is more solvent-exposed than the other in an HlyB dimer. The side-view of the SAXS 3D shape reconstruction (**Figure 2E**, right) reveals density “above” the CLD that reaches into the membrane region. While parts of this density may originate from membrane lipids retained from the purification process of the stalled complex or bound GDN detergent, we observe motions of the HlyA2-bound CLD2 from HlyB_I_ in the MD simulations that can also account for parts of this density. CLD2 thereby reaches the headgroup region of the lower membrane leaflet.

We analyzed the MD simulations of both systems for potential additional binding sites of HlyA2 on HlyB (**Figure 2F**). Besides interactions with CLD2, interactions with the lateral gate entrance of the HlyB dimer were observed frequently in both systems. In addition, HlyA2 interacted with the NBD2 domain in the HlyA2/HlyB_OOO_/HlyD system (NBD2 is in the same protomer as CLD2). Several of the observed interaction sites (Y477, F518, Y519, V682, L697, Y700) were previously identified to be critical for the interaction with the toxin (33). This result was unexpected because in this previous study (33) HlyA1 or HlyA were used, which both contain the secretion signal, which was implicated to be important for the interaction with the NBD. In turn, no binding between HlyA2 and an isolated NBD was observed (33). The results from our simulations thus lead to the suggestion that a high local concentration of HlyA2 due to its interaction with the CLD2 and the lateral gate entrance may also favor interactions with the NBD, despite the lack of the secretion signal.

### HlyA2 binding to HlyB_IOO_/HlyD can stabilize neighboring HlyD particularly in those regions that are critical for recruiting TolC

To assess the influence of HlyA2 binding on the structural stability of the HlyB/D complex, we used conformations generated from unbiased MD simulations and applied a dynamic allostery model based on Constraint Network Analysis (24, 26, 41) previously used on bacterial transporters (42). This model describes allosteric effects in terms of the per-residue (*i*, eq. 1, eq. 2) or per-region (*r*, eq. 3) rigidity index Δ*r*_i/r,(bound-apo)_ (41). We evaluated Δ*r*_i/r,(bound-apo)_ for HlyB1, HlyB2, HlyD1, and HlyD2 in (HlyA2)/HlyB_OOO_/HlyD and (HlyA2)/HlyB_IOO_/HlyD; HlyB1 contains CLD1 and HlyB2 the CLD2 in the bound dimer, and HlyD1 and HlyD2 are the respective neighboring HlyD chains.

The dynamic allostery model reveals a configuration– and toxin binding-dependent impact on the structural stability of HlyB1, HlyB2, HlyD1, and HlyD2 (**Figure 3A**). The structural stability of HlyB_OOO_/HlyD is not markedly affected by toxin binding, in contrast to changes observed in HlyB_IOO_/HlyD. The binding of HlyA2 destabilizes the HlyB_I_ dimer, while HlyD1 and HlyD2 become stabilized. The largest stabilizing effects occur at a charged motif (R_34_EKDE_38_) in the cytoplasmic domain, located at the HlyD N-terminus forming contacts with the HlyB monomer, previously identified as essential for TolC recruitment (17) (**Figure 3B**). This motif is at a distance of ∼50 Å from the CLD2 domain. The stabilizing effect also reaches the top region of HlyD at a distance of ∼250 Å from the CLD2. Together, this reveals a long-range effect of HlyA2 binding that percolates through the structure. By contrast, only small changes are observed in the HlyB_O_ dimers and their neighboring HlyD protomers.

**Figure 3.**
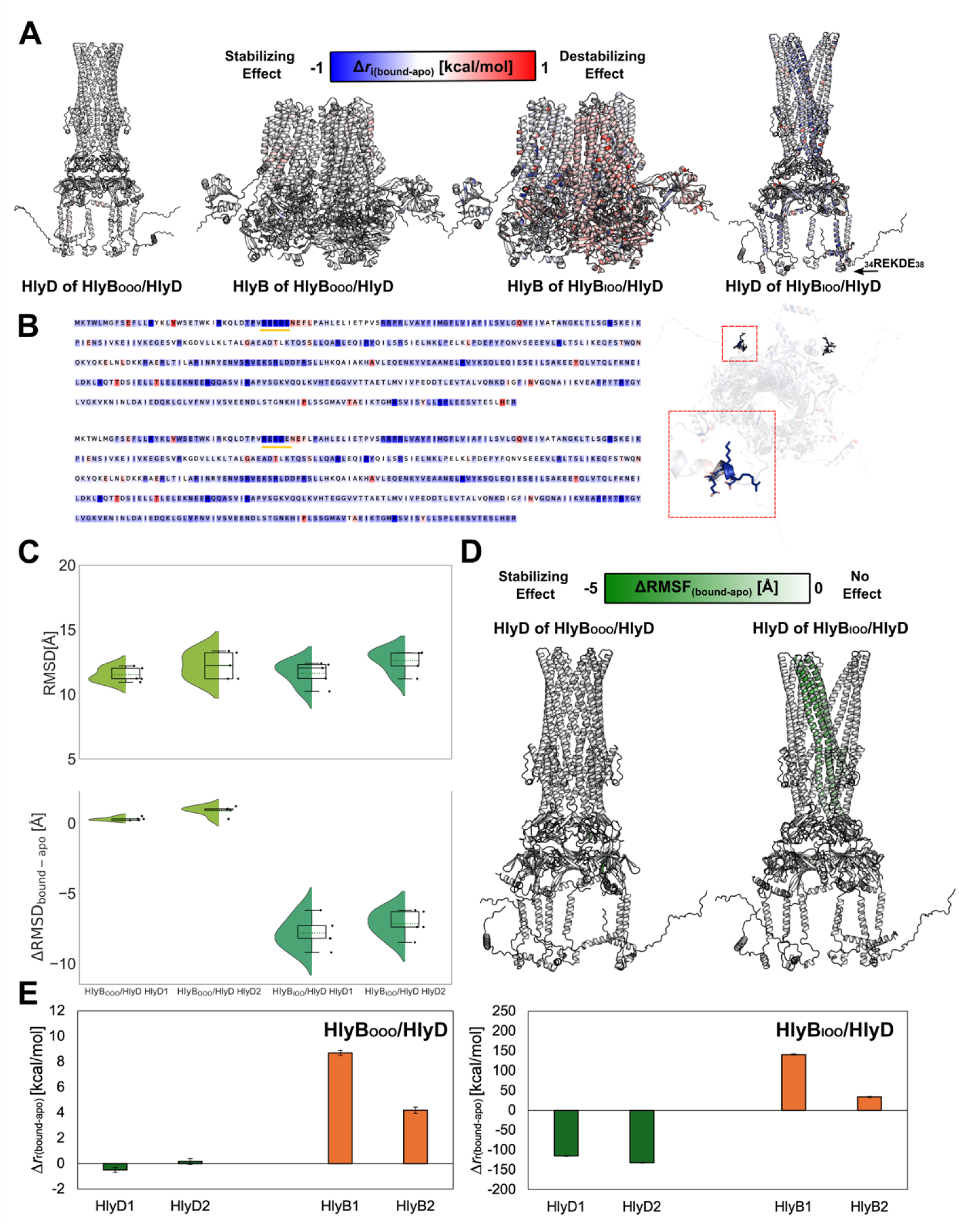
Long-range effects in HlyB/HlyD systems due to HlyA2 binding. **A)** Δ*r*_i,(bound-apo)_ (eq. 2) obtained from rigidity analysis of MD ensembles mapped at the residue level onto the average structure of HlyB_OOO_/HlyD (left) and HlyB_IOO_/HlyD (on the right). For clarity, the HlyB and HlyD components are depicted separately. Red (blue) secondary structure color indicates a destabilizing (stabilizing) effect after HlyA2 binding. The _34_REKDE_38_ motif is also indicated with a black arrow. **B)** The changes in Δ*r*_i,(bound-apo)_ were also mapped at the sequence level for HlyD1 (top) and HlyD2 (bottom) in HlyB_IOO_/HlyD. In both cases, the highest stabilizing contribution was found for the _34_REKDE_38_ motif (highlighted sequence on the left, and shown as zoomed cartoon from an intracellular view of HlyD on the right). **C)** RMSD of HlyD1 and HlyD2 in HlyB_OOO_/HlyD and HlyB_IOO_/HlyD (*top*) and ΔRMSD_bound-apo_ of HlyD1 and HlyD2 in HlyA2/HlyB/HlyD versus HlyB/HlyD (*bottom*) plots. The violin plots indicate the distribution of the data points; the inner box of a box plot represents the interquartile range, with the horizontal black line indicating the median. The dotted green line represents the mean, and the whiskers show the rest of the distribution, excluding points determined to be outliers when they fall outside 1.5 times the interquartile range. The light green violins indicate the (HlyA2)/HlyB_OOO_/HlyD system and the dark green violins the (HlyA2)/HlyB_IOO_/HlyD system. **D)** ΔRMSF_bound-apo_ mapped at the residue level on HlyD of (HlyA2)/HlyB_OOO_/HlyD (left) and (HlyA2)/HlyB_IOO_/HlyD (right). Darker green motifs indicate reduced structural variability of such residues after HlyA2 binding. **E)** Cumulative effect on the structural stability (Δ*r*_r,(bound-apo)_, eq. 3) for HlyD1 and HlyD2 (green histogram bars), HlyB1 and HlyB2 (orange) of (HlyA2)/HlyB_IOO_/HlyD (left) and (HlyA2)/HlyB_IOO_/HlyD (right). The SEMs for *n* = 5 replicas are also shown.

The stabilizing effect on HlyD1 and HlyD2 is mirrored in the structural variability of these protomers: In the apo state, the structural variabilities of these protomers in HlyB_OOO_/HlyD and HlyB_IOO_/HlyD are similar, with the HlyD2 protomers showing moderately higher values (**Figure 3C**, top). Binding of HlyA2 reduces the structural variability of the neighboring HlyD1 and HlyD2 considerably in the case of HlyA2/HlyB_IOO_/HlyD but has a negligible impact in the case of HlyA2/HlyB_OOO_/HlyD (**Figure 3C**, bottom). On a per-residue level, this impact is also seen in a pronounced reduction of atomic positional fluctuations around the average position for HlyA2/HlyB_IOO_/HlyD but not for HlyA2/HlyB_OOO_/HlyD (**Figure 3D**). Large decreases were found for those regions of HlyD1 and HlyD2 that are involved in TolC binding (**Figure S8**). These are up to 250 Å away from the CLD2. Finally, summing the Δ*r*_i,(bound-apo)_ over HlyB1, HlyB2, HlyD1, and HlyD2 yields an overall effect for the region, Δ*r*_r,(bound-apo)_, which corroborates that HlyA2 binding impacts HlyB_IOO_/HlyD markedly more than HlyB_OOO_/HlyD and that a pronounced stabilization of HlyD1 and HlyD2 occurs only in the former case (**Figure 3E**). These findings suggest that HlyA2 binding to HlyB_IOO_/HlyD can stabilize neighboring HlyD, particularly in those regions that are critical for recruiting TolC.

### Long-range effects due to HlyA2 binding extend to TolC

To investigate the involvement of stabilized HlyD domains in TolC interaction, we generated the, to our knowledge, first full structural model of the HlyB/HlyD/TolC complex (**Figure 4A**) using a fragmented modeling approach, which overcame GPU memory limitations for large protein assemblies. To validate the model, SAXS on the stalled eGFP-HlyA/HlyB/HlyD/TolC complex was used to check whether the T1SS complex was fully assembled and whether the computational structural model matched the calculated ab initio model. The molecular weight of the T1SS complex was determined to be 1086 kDa, which is in agreement with the theoretical molecular weight of 1106 kDa (without detergent) (Table S3). The spatial dimensions of the theoretical model show an *R_g_* of 12.00 nm and *D_max_* of 44.01 nm, which is in good agreement with the experimentally determined *R_g_* of 12.17 nm and *D_max_* of 46.20 nm, respectively. The calculated ab initio model superimposed onto the computational structural model is shown in **Figure 4A**.

**Figure 4.**
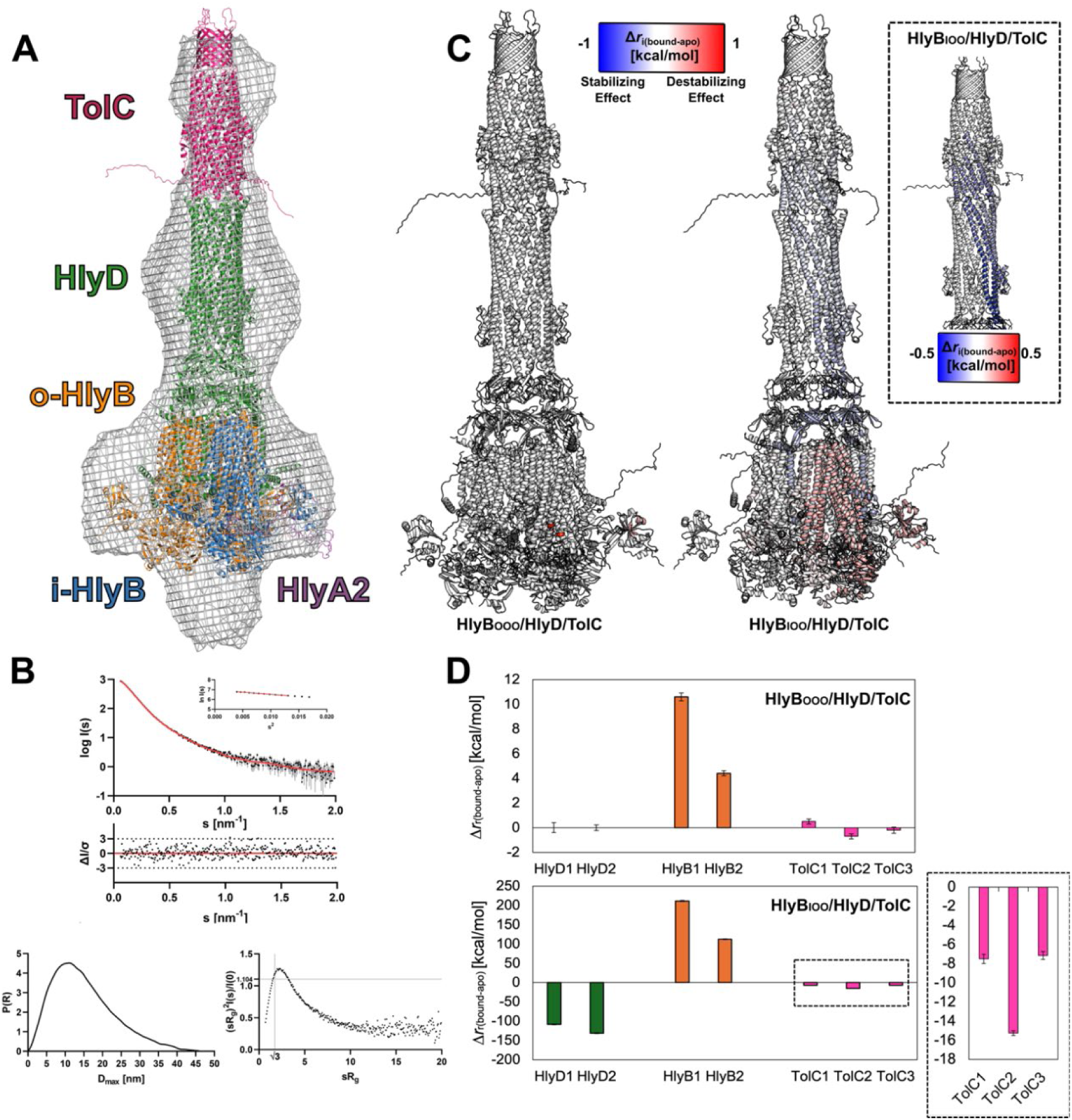
HlyA2/HlyB_IOO_/HlyD/TolC complex model shows agreement with SAXS 3D shape reconstruction. **A)** *Top:* The generated complex model superimposed on the calculated SAXS 3D shape reconstruction (*top*). The full density is shown as a grey mesh; HlyA2, HlyB_IOO_, HlyD, and TolC are shown as purple, blue (HlyB_I_), orange (HlyB_O_), green, and magenta ribbons, respectively. *Bottom:* **B)** *Top:* Experimental scattering data of T1SS. Experimental data are shown in black dots, with grey error bars. The DAMMIF ab initio model fit is shown as red line; below is the residual plot of the data. The Guinier plot of T1SS is added in the right corner. *Bottom-left: p(r)* function of T1SS. *Bottom-right:* Dimensionless Kratky plots of T1SS. **C)** Δ*r*_i,(bound-apo)_ (eq. 2) mapped at the structural level for (HlyA2)/HlyB_OOO_/HlyD/TolC (left) and (HlyA2)/HlyB_IOO_/HlyD/TolC (right). Red (blue) secondary structure color indicates a destabilizing (stabilizing) effect after HlyA2 binding. The blow-up on the right focuses on HlyD and TolC. **D)** Cumulative effect on the structural stability (Δ*r*_r,(bound-apo)_, eq. 3) on neighboring HlyD1 and HlyD2 (green histogram bars), HlyB1 and HlyB2 (orange) and TolC1, TolC2, and TolC3 (magenta) for (HlyA2)/HlyB_OOO_/HlyD/TolC (*top*) and (HlyA2)/HlyB_OOO_/HlyD/TolC (*bottom*) configurations. The blow-up on the right focuses on TolC. The SEMs for *n* = 5 replicas are shown.

Four transporter states were modeled ((HlyA2)/HlyB_OOO_/HlyD/TolC and (HlyA2)/HlyB_IOO_/HlyD/TolC, each one apo or HlyA2-bound to the CLD2 of one of the HlyB_O_ dimers or to the CLD2 of HlyB_I_). These structures were subjected to the model of dynamic allostery, revealing patterns of structural stabilization in HlyD1 and HlyD2 and structural destabilization in HlyB_I_ of (HlyA2)/HlyB_IOO_/HlyD/TolC (**Figure 4B**), consistent with the results for (HlyA2)/HlyB_IOO_/HlyD (**Figure 3A**). Likewise, and again consistently, almost no influence on structural stability is found for (HlyA2)/HlyB_OOO_/HlyD/TolC. Note that here the analysis of long-range effects was performed using an ensemble of network topologies generated with the fuzzy network constraints approach (43). Interestingly, particularly in (HlyA2)/HlyB_IOO_/HlyD/TolC, the stabilizing effect extended from HlyD to TolC (**Figure 4B, C**). Residues D371 and D374 of TolC, which were predicted to be important for tunnel constriction (21), were allosterically impacted in a pronounced extent (Δ*r*_i,(bound-apo)_ ∼ –0.20 to –0.25 kcal/mol). A similar threshold was previously applied to detect marked stability changes in this context (44, 45). This confirms that allosteric signaling through HlyD is necessary for TolC recruitment and activation (22, 46) and may help explain proposed mechanisms where substrate binding induces conformational changes that open TolC for efflux (17–19).

## Discussion

Considering that many mechanistic details of HlyA translocation through T1SS have remained unclear, in this study, we addressed early steps of the transport process combining structural modeling with atomistic MD simulations and biochemical and SAXS experiments. Our data indicate that CLD1 is structurally the least variable in HlyB_OOO_/HlyD and HlyB_IOO_/HlyD, and CLD2, described to be functionally relevant for toxin translocation (15, 33), is so in HlyB_IOO_/HlyD, that HlyA2 binding to HlyB_IOO_/HlyD can stabilize neighboring HlyD particularly in those regions that are critical for recruiting TolC, and that long-range stabilizing effects due to HlyA2 binding extend to TolC.

Initially, we generated complete models of HlyB/HlyD complexes with distinct combinations of HlyB in the inward-facing or occluded conformation. Besides complementing the missing parts in the two cryo-EM structures HlyB_OOO_/HlyD and HlyB_IOO_/HlyD, we also generated complexes with HlyB_IIO_/HlyD and HlyB_III_/HlyD configuration. In doing so, we aimed at using as much as possible structural knowledge from experimental data by following a fragmented modeling approach. Overall, the obtained complex models are of very good quality as indicated by a comparative analysis of Ramachandran statistics across obtained systems (Figure S2). Still, as consistently found in MD replicas, HlyB_OOO_/HlyD and HlyB_IOO_/HlyD showed the least structural variability in functionally relevant parts. Only HlyB_OOO_/HlyD and HlyB_IOO_/HlyD configurations have been found in cryo-EM experiments (15). Together, this suggests that the differences in structural variability arise from the increased prevalence of HlyB_I_ in HlyB_IIO_/HlyD and HlyB_III_/HlyD, rather than from a differing quality of the starting structures.

We used HlyB_OOO_/HlyD and HlyB_IOO_/HlyD to scrutinize the specific roles of the CLD2 of HlyB. This domain, previously predicted to be important for substrate recognition, has not been fully resolved in recent cryo-EM maps (15). Our work also introduced the first HlyA2-bound HlyB_OOO_/HlyD and HlyB_IOO_/HlyD models by overcoming structural prediction limitations of AlphaFold3 (36), which incorrectly generated a structured HlyA2 at the CLD interface, although HlyA2 is known to interact with the CLD in an unfolded state (23). MD simulations revealed only for HlyB_OOO_/HlyD additional interactions of HlyA2 with the NBD domain of the HlyB protomer that also interacts with HlyA2 via its CLD2. The HlyA2-NBD interactions involved residues typically interacting with the secretion signal sequence, which is missing in HlyA2 (33, 47). Thus, it cannot be excluded that the interaction between the RTX motif of HlyA2 and the CLD, which was missing in these experiments but is present in our simulations, has an impact on the interaction with the NBD. By contrast, in the HlyA2/HlyB_IOO_/HlyD complex, no interaction with the NBD was found, and the HlyA2-bound CLD2 showed an increased structural variability compared to the apo CLD2. The increase in variability may be important to help shuttle the C-terminus of HlyA into the HlyB_I_ dimer pore for transport. The motions of the HlyA2-bound CLD2 from HlyB_I_ observed in the MD simulations could account for parts of the SAXS density of the HlyA/HlyB/HlyD/TolC complex observed “above” the CLD. In turn, apo CLD2 of HlyB_O_ in HlyB_OOO_/HlyD by itself is rather variable, so it may act as an “encounter element” for binding HlyA. Mobile domains or even disordered structural parts have been described before to act as encounter elements in other biomolecular systems (48, 49). Together, the data allows one to speculate that HlyB_OOO_/HlyD is the initial configuration to which HlyA binds, supported by a high structural variability of apo CLD and the possibility to interact with the NBD. The bound HlyB_O_ is suggested to transition to an HlyB_I_ conformation, which might foster the insertion of HlyA into the HlyB transport pathway (Figure 5). Due to the limited simulation length, we could not observe this transition in our MD simulations, however.

**Figure 5.**
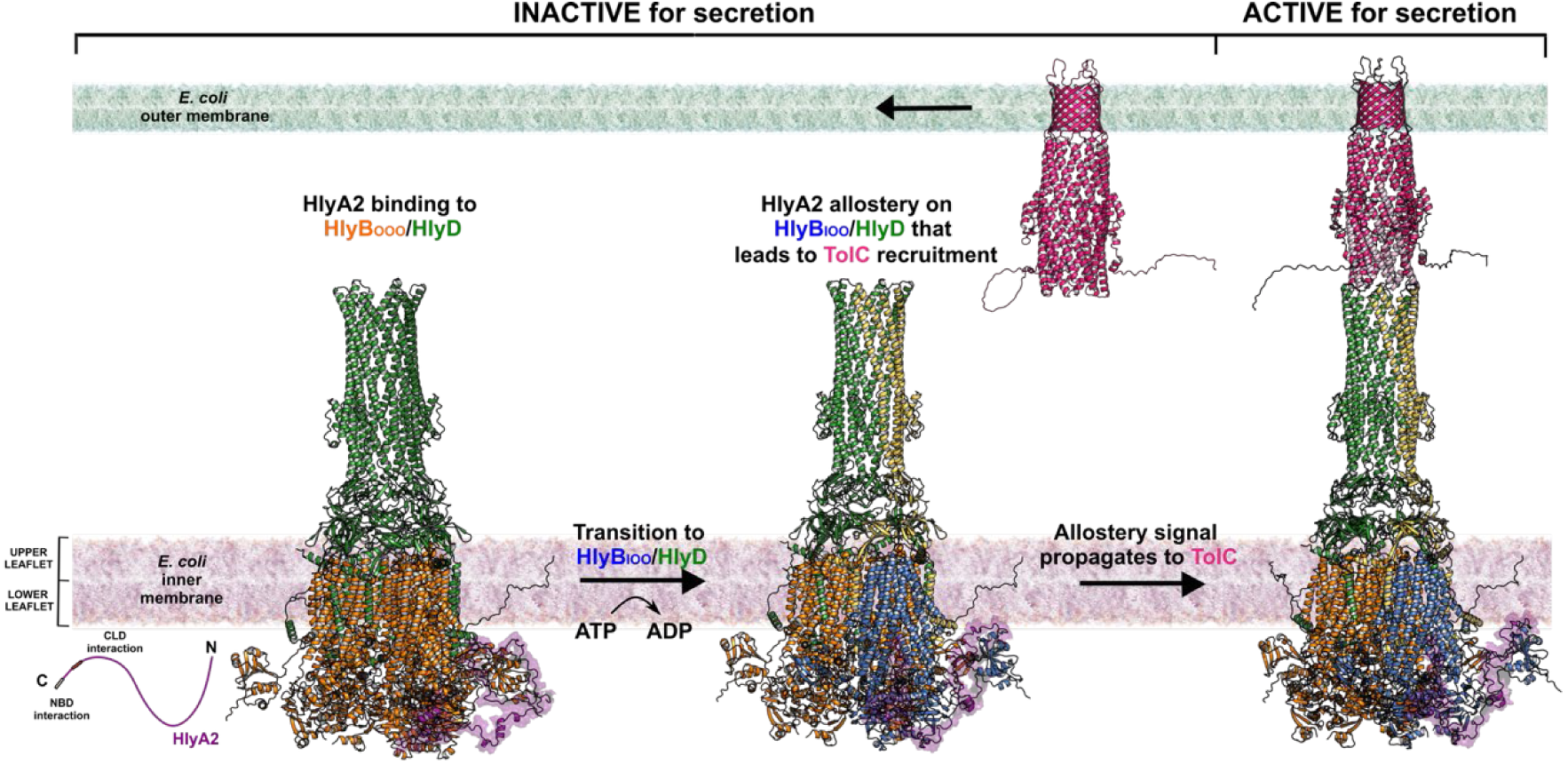
A working model illustrating the potential impact of HlyA on HlyB/HlyD/TolC. The HlyB/HlyD complex (IMC) is anchored to the inner membrane through HlyB (orange) and parts of HlyD (green). HlyA2 (purple) corresponds to the C-terminal fragment of HlyA (residues 807-966), containing the three conserved RTX repeats but being devoid of the last 57 amino acid residues and, therefore, the secretion signal. According to the proposed model, HlyA2 binds to the HlyB_OOO_/HlyD configuration. After the transition to a HlyB_IOO_ configuration (HlyB_I_ is colored blue), an allosteric signal propagates through the HlyD chains neighboring the inward-facing dimer (HlyD1 and HlyD2, in yellow). We suggest that the allosteric signal may contribute to recruiting TolC (magenta). In the complex model with TolC, the allosteric signal propagates into TolC. Secretion of HlyA might then occur through a single-inward-facing dimer.

Next, we applied a model of dynamic allostery (26, 41, 43) based on rigidity theory (24, 45) to (HlyA2)/HlyB_OOO_/HlyD and (HlyA2)/HlyB_IOO_/HlyD to probe for long-range effects due to HlyA2 binding. The model has been used before on a variety of biological systems, including enzymes (50), ion channels (44), and transporters (42). The analysis highlighted the role of HlyB_I_ in TolC recruitment after HlyA2 binding. Specifically, the REKDE motif of HlyD, implicated in TolC interaction (17), exhibited the largest stabilization upon HlyA2 binding. The allosteric effect was most pronounced in the HlyD protomers adjacent to HlyB_I_. Given that all three dimers are capable of ATP hydrolysis and substrate transport (15), the allosteric effect may subsequently occur to all HlyD chains neighboring the respective inward-facing HlyB dimer. The increased structural stability in HlyD was supported by a decreased structural variability observed in MD simulations for the HlyA2-bound state of HlyB_I_ compared to the apo state. The increased structural stability of HlyD observed upon HlyA2 binding may also explain why the published cryo-EM maps of the apo HlyB/HlyD complex lack density in the upper region of HlyD and supports the substrate-induced assembly mechanism proposed for HlyD (15).

Finally, we generated the first atomistic structural model of the HlyA2/HlyB/HlyD/TolC complex, considering the established stoichiometry (15). The model fits into the SAXS density obtained for a stalled HlyA/HlyB/HlyD/TolC complex. Applying our model of dynamic allostery, we observed a stabilizing effect at the HlyD/TolC interface, particularly involving residues D371–D374, which are known to be important for TolC tunnel constriction (21). Notably, this stabilization occurred throughout the entire TolC although the strongest effects were found for that part of TolC that is closest to the HlyD chains neighboring HlyB_I_. These findings suggest that the interaction of HlyA with a single CLD2 may be sufficient to recruit TolC.

In summary, this study provides new molecular insights into the role of the CLD domain of *E. coli* T1SS in HlyA2 substrate recognition and indicates how substrate recognition can lead to allosteric signal transmission over a distance up to 250 Å, which may facilitate the assembly of the full T1SS complex. The full T1SS model provides a foundation for future molecular simulation studies of T1SS. A deeper understanding of hemolysin transport could inform strategies to inhibit this secretion machinery, presenting a promising avenue for the development of novel antibiotic strategies targeting *E. coli*.

## Materials and Methods

### Construction of HlyB/HlyD and HlyB/HlyD/TolC models in the apo– and HlyA2-bound configuration using MSAs– and template-based information

Modeling of a fully multimeric HlyB/HlyD (3×2: 6) protein complex structure as a single entity in ColabFold (28) was not technically possible at the time of generating the models due to GPU memory constraints. To overcome this, a fragmented modeling approach was implemented, in which different regions of the complex were modeled independently and later merged by overlapping regions, with gaps filled using MODELLER (51). The following structures were used: 1) the cryo-EM HlyB/HlyD complex (PDB ID: 7SGR; configuration single-inward) as the central scaffold (15), 2) a 2xHlyB protomer/2xHlyD protomer ColabFold model (configuration fully-occluded) (**Figure S1A**), 3) a 6xHlyD protomer ColabFold model (**Figure S1B**). To obtain an outward-occluded structure and maintain consistency, dimer B from PDB ID 7SGR was copied and overlayed onto dimers A and C using PyMOL (52). The 2xHlyB/2xHlyD model was then overlayed and the missing CLD was copied and positioned, avoiding steric clashes with neighboring protomers. HlyD regions present in PDB ID 7SGR were retained as references for placing the matching regions from the 6xHlyD model. The assembled structures were loaded into MODELLER (51), using for every other HlyD chain regions 82-458 or 78-459 from the 6xHlyD model, and regions 29-81 and 459-474 or 9-77 and 460-478 from PDB ID 7SGR (14), respectively. Missing loops were filled by MODELLER (51). The adenosine triphosphate (ATP) bound to *E. coli* HlyB_O_ together with the coordinating Mg^2+^ is available in the Protein Data Bank (PDB ID 8DCK) (15). After superimposing the experimental protein structures with our generated model (RMSD = 0.9 Å), the bound poses of ATP and Mg^2+^ were extracted and merged with the predicted model. In parallel, ATP was docked into the single protomers, using the same protocol we used already for other bacterial membrane proteins (42). From the fully-occluded model (HlyB_OOO_/HlyD), “inward-facing” configurations with one (HlyB_IOO_/HlyD), two (HlyB_IIO_/HlyD), or three dimers (HlyB_III_/HlyD) were generated (Table S1). Each inward-facing HlyB dimer was created by superimposing PDB ID 7SGR (15) onto the fully-occluded model, replacing overlapping regions with chains B and C from PDB ID 7SGR, and positioning the missing CLD to avoid overlap with neighboring protomers. The structural quality of the starting conformations was assessed in terms of Ramachandran distributions calculated with MolProbity (53) (**Figure S2**).

For HlyA2-bound systems (HlyA2/HlyB_OOO_/HlyD and HlyA2/HlyB_IOO_/HlyD; Table S1), a structural model of the unfolded toxin at ambient temperature was required. HlyA and other RTX toxins feature glycine– and aspartate-rich nonapeptide repeats, known as GG repeats, which bind Ca^2+^ and trigger folding, after secretion (23). While HlyA2 has been shown to fold in the presence of Ca^2+^, it does not do so intracellularly or at CLD domains (23). For these complexes, HlyA2 (residues 807-966), a C-terminal fragment containing three RTX repeats but lacking the secretion signal, was used. Residues on the CLD interacting with HlyA2 were identified from NMR CSP and mutagenesis studies (33). Since the most recent structure prediction tools failed to model HlyA2 in an unfolded state (**Figure S3**), a stepwise approach was used. Initially, the GG repeat motifs were modeled in a linear unfolded state using Peptide Discovery from the Schrödinger software suite (39). Then, peptide docking of the GG repeats motifs to the CLD domain was performed constraining the H-bond interactions known from the literature using the Glide suite from Schrödinger (37, 38, 54). Afterward, the remaining part of the peptide was designed using the 3D Builder option for each residue, and a minimization step was performed for each introduction. In this way, a HlyA2/CLD complex (Table S1) was generated based on the published interactions.

As only one of the two CLD domains (CLD2) is solvent exposed and competent to bind HlyA in the cytoplasm, we then superimposed the HlyA2/CLD model with the respective HlyB/HlyD model by superimposing the CLD2 and subsequently merging the bound HlyA2 (RMSD = 0.5 Å) to obtain a full occluded HlyB-D complex carrying the toxin (HlyA2/HlyB_OOO_/HlyD) and single inward-facing HlyB-D complex carrying the toxin on the inward-facing HlyB_I_ (HlyA2/HlyB_IOO_/HlyD).

The generated HlyB/HlyD models were extended to construct HlyB/HlyD/TolC models in the presence and absence of bound HlyA2. To design the interaction between HlyD and TolC, a 6xHlyD(residues 132-385)/3xTolC ColabFold model was generated (**Figure S1C**). TolC was positioned in the HlyB/HlyD complex by superimposing the 6xHlyD(residues 132-385)/3xTolC model with the corresponding region in the 6xHlyD model, and partial HlyD structures were removed.

### Preparation of the starting structures for molecular dynamics simulations of HlyB/HlyD

The generated HlyB/HlyD models were used to perform molecular dynamics (MD) simulations for the systems listed in Table S1, except HlyA2/CLD. All models were oriented in the membrane using the PPM server (55). Starting configurations were embedded into a DOPE/DOPG = 3:1 membrane, matching the native inner membrane composition of Gram-negative bacteria (31), and solvated using PACKMOL-Memgen (56, 57). ATP was parametrized using a protocol described in the Supporting Text. Crosslinking results indicated that the substrate HlyA is translocated through a HlyB dimer and not through the central pore, which is partially filled with phospholipids in the cryo-EM structures (PDB IDs 7SGR, 8DCK) (15). We thus filled the central pore in HlyB/HlyD complexes with phospholipids as well. To calculate the amount of lipids required, we used the following protocol. In preliminary simulations, we observed a significant penetration of water into the central pore of HlyB. We defined the pore region as the intersection of a spherical region with a radius of 30 Å localized at the protein’s center of mass and a vertical double-sided range of 34 Å length around the center of the membrane. Within this volume, we detected 795 water atoms in the lower part and 276 water atoms in the upper part, starting from the protein’s center of mass. Considering a volume of 30 Å^3^ per water molecule and three atoms per water molecule, the volume of a region to be filled with lipids amounts to

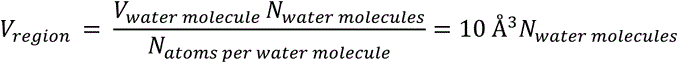

Thus,

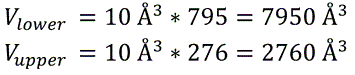

Using an estimation of about 1300 Å^3^ per lipid, we have placed 2 DOPG molecules in the lower part and 6 DOPE molecules in the upper part of the pore, respectively, and kept them restrained using the leaflet restraint available in PACKMOL-Memgen and the option “inside cylinder” placement (57). A distance of at least 15 Å between the protein or membrane and the solvent box boundaries was kept. To obtain a neutral system, counterions were added that replaced solvent molecules (0.15 M KCl).

The HlyA2/CLD model was also used to perform MD simulations. The solvation and parametrization protocols used are similar to those for the (HlyA2)/HlyB/HlyD systems. The generated systems are summarized in Table S1. The size of the resulting systems was ∼2*10^5^ atoms for the system CLD and ∼2*10^6^ for the remaining systems.

### Unbiased molecular dynamics simulations

The GPU implementation of Particle Mesh Ewald MD simulations (pmemd) from the AMBER22 suite of molecular simulation programs (58) with the ff19SB (59) and Lipid 21 (60) forcefields for the protein and membrane lipids, respectively, were used; water molecules and ions were parametrized using the OPC3POL model (61, 62) and the Li and Merz 12-6 ions parameters (63, 64). For each configuration in Table S1, five independent MD simulations of 1 μs length were performed. Covalent bonds to hydrogens were constrained with the SHAKE algorithm (65), and the hydrogen masses were repartitioned (66), allowing the use of a time step of 4 fs. Details of the thermalization of the simulation systems are given below. All unbiased MD simulations showed structurally rather invariant protein structures and membrane phases as evidenced by electron density calculations (**Figure S5**). The overall RMSD of ATP-Mg^2+^ revealed structurally invariant binding poses across different replicas except that for the full-inward-facing complex system (**Figure S6**).

### Relaxation, thermalization, and production runs of the obtained systems

An initial energy minimization step was performed with the CPU code of pmemd (67). Each minimization was organized in four steps of 5000 cycles each, for a total of 20000 cycles of steepest descent minimization. Afterward, each minimized system was thermalized in one stage from 0 to 300 K over 125 ps using the NVT ensemble and the Langevin thermostat (68), and the density was adapted to 1.0 g cm^-3^ over 500 ps using the NPT ensemble with a semi-isotropic Berendsen barostat (69) with the pressure set to 1 bar. The thermalization and equilibration were performed with the GPU code of pmemd (67). There were ten density equilibration steps with a total time of 19375 ps. The sum of the thermalization, density adaptation, and equilibration took 20 ns.

For each replica, 1 μs of production run using the GPU code of pmemd was performed in the NPT ensemble at a temperature of 300 K using the Langevin thermostat (68) and a collision frequency of 1 ps^-1^. To avoid noticeable distortions in the simulation box size, semi-isotropic pressure scaling using the Berendsen barostat (69) and a pressure relaxation time of 1 ps was employed by coupling the box size changes along the membrane plane (70).

### Analysis of MD trajectories of the prepared systems

The trajectories were analyzed with CPPTRAJ (71, 72). Root-mean-square-deviation (RMSD) calculations were performed to describe the structural variability of the different protein complexes. From the obtained trajectories, we calculated the 3D density maps of the CLD domains considering all atoms using the grid function available in CPPTRAJ with a grid spacing of 1.5 Å. We applied a contour level of 1σ (one standard deviation above the mean value) as already done for other bacterial membrane proteins (32).

### Constraint network analysis of the HlyB/HlyD ensembles

To detect changes in structural rigidity between HlyB/HlyD (full-occluded and single-inward complexes) in the apo and HlyA2-bound structures, we analyzed ensembles of constraint network topologies based on conformational ensembles saved every 400 ps from the production phase of the unbiased MD simulations. For each configuration, we generated 400 conformations. Overall, we investigated 8,000 conformations in this study.

Structural rigidity was analyzed with the CNA software package (24, 26), which is a front and back end to the FIRST software (25). It was used to construct networks of nodes (atoms) and covalent and noncovalent (hydrogen bonds, salt bridges, and hydrophobic tethers) constraints. The hydrogen bond energy (including salt bridges) is determined from an empirical function (74), while hydrophobic tethers between carbon and sulfur atoms were considered if the distance between these atoms was less than the sum of their van der Waals radii plus a cutoff of 0.25 Å (27, 75). Biomolecules generally display a hierarchy of rigidity that reflects the modular structure of biomolecules in terms of secondary, tertiary, and super-tertiary structure (24). This hierarchy can be identified by gradually removing noncovalent constraints from an initial network representation of a biomolecule, which generates a succession of network states *σ*. Only those hydrogen bonds were kept that have an energy *E_HB_* ≤ *E_cut_*(*σ*). In our analysis, we used energy cutoffs between *E_start_* = –0.1 kcal mol^-1^ and *E_stop_* = –10.0 kcal mol^-1^ with a step size of *E_step_* = 0.1 kcal mol^-1^. We also assigned the cutoff for finding native contacts between pairs of residues (ncd = 4.5). For all MD-generated snapshots, we calculated a rigidity index *r_i_* (41) (eq. 1).

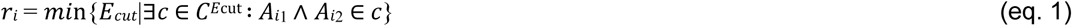

The index is defined for each covalent bond *i* between two atoms *A_i_*, as the *E_cut_* value during a constraint dilution simulation at which the bond changes from rigid to flexible. Phrased differently, this index monitors when a bond segregates from any rigid cluster *c* of the set of rigid clusters *C*^*E*cut^. It reflects structural stability on a per-residue basis and, thus, can be used to identify the location and distribution of structurally weak or strong parts in the protein complex.

The same analysis was performed for the five independent replicas of (HlyA2)/HlyB_OOO_/HlyD and (HlyA2)/HlyB_IOO_/HlyD, either in the apo or HlyA2-bound form. The per-residue *r*_*i*_ values were averaged on a per-residue basis across the five independent replicas and the SEM was calculated. Afterward, the per-residue difference Δ*r*_i,(bound-apo)_ (eq. 2) was calculated, including an error propagation analysis for the SEM. Values of Δ*r*_i,(bound-apo)_ < 0 (> 0) indicate that the bound structure is more rigid (flexible) than the apo structure.

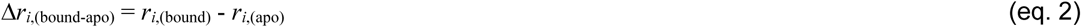

Finally, we obtained the cumulative difference Δ*r*_r_ of a specific region of interest, *r*_r_ (HlyB or HlyD) (eq. 3) by summing the *m* single per-residue Δ*r*_i,(bound-apo)_ contributions, including an error propagation analysis.

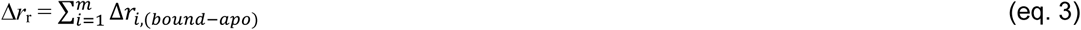

The error propagation for the SEM was calculated for both eq. 2 and eq. 3 using eq. 4 and eq. 5, respectively.

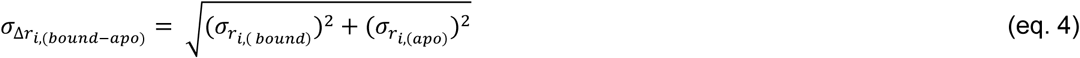

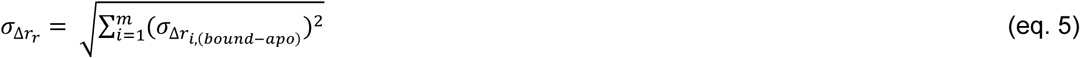

For the HlyB/HlyD/TolC systems, thermal unfolding simulations were performed using the ENT^FNC^ approach (via the –-ent_fnc flag in CNA). The ENT^FNC^ approach performs rigidity analyses on ensemble network topologies (ENT) generated from a single input structure. The ENT is based on definitions of fuzzy noncovalent constraints (FNC) derived from persistency data of noncovalent interactions from MD simulations. Therefore, the approach considers thermal fluctuations of a biomolecule without sampling conformations (24). The Δ*r*_i,(bound-apo)_ and the Δ*r*_r_ calculations follow the same scheme as for the MD-generated unfolding simulations.

### Cloning

The generation of the pK184 plasmid encoding HlyB and HlyD was described previously here (34). A plasmid containing HlyA with 6xHis followed by eGFP tag at the N-terminus (pSOI-6xHis-eGFP-HlyA) (40) was amplified, and overhangs were generated at position L714 in *hlyA* gene. Sequences of three copies of the FLAG tag separated each by a GGGGS linker (FLAGx3-L) in the form of two single-stranded oligos (Eurofins) were mixed for hybridization. The FLAGx3-L sequence was inserted into the *hlyA* gene using Gibson Assembly (NEB) resulting in the plasmid pSOI-6xHis-eGFP-HlyA-Flagx3-L. All oligonucleotides used in this study are summarized in Table S2.

### Expression

For co-expression of the stalled complex, *E. coli* BL21(DE3) competent cells were co-transformed with both the plasmids and grown on LB agar plates supplemented with 100 μg·ml^−1^ ampicillin and 30 μg·ml^−1^ kanamycin. Overnight cultures using colonies (bearing the double plasmids) were used to inoculate 25 ml of LB medium supplemented with both antibiotics at an optical density at 600 nm (OD_600_) of 0.1. The cultures were grown at 37°C and 180 rpm.

Initially, the expression of HlyB, and HlyD was induced at an OD_600_ of 0.8 to 1.0 using 1 mM IPTG and the cells were grown for half an hour. CaCl_2_ was added to the medium at a final concentration of 5 mM and the expression of HlyA was induced using 10 mM L-arabinose. Cells were further expressed for 1 h. The cell pellet was collected by centrifugation (4500 rpm, 20 min), resuspended in buffer A (50 mM Tris-Cl pH 7.5, 50 mM KCl, 2 M NaCl, 10 mM CaCl_2_).

### Purification of the stalled complex

The cell pellet was lysed by one pass through a high-pressure homogenizer at 1000 bar (Microfluidics Cell disruptor). Cell lysate was centrifuged at 10,000 x g for 30 min, and the supernatant was subjected to a second round of ultracentrifugation at 1000,000 x g for 4 hours to pellet the cell membrane. After being dispersed with a hand-held homogenizer in buffer A, the membrane fraction was solubilized with buffer B (30 mM Tris pH 8.0, 150 mM NaCl, 10 mM CaCl2, 15 % glycerol) plus 1% GDN (Anatrace) and two complete EDTA-free protease inhibitor tablets (Roche) for overnight at 4°C. The insoluble fraction was removed by centrifugation at 35800 rpm for 30 min (Beckman type 45 Ti rotor). To purify the complex, the supernatant was first diluted 1:10 using buffer C (50 mM Tris-Cl pH 7.5, 150 mM NaCl, 10 mM CaCl2) and applied to ANTI-FLAG® M2 Affinity Gel (Sigma Aldrich) multiple times. The resin was washed with 20 column volumes of buffer D (50 mM Tris-Cl pH 7.5, 150 mM NaCl, 10 mM CaCl2 and 0.0063% GDN), and the protein was eluted with buffer D plus 150 ng/μl 3xFLAG® peptide (Pierce^TM^ 3x DYKDDDDK Peptide; ThermoFisher Scientific). The eluate was then concentrated (100 kD cut-off Eppendorf concentrator, Amicon) and further purified by size exclusion chromatography (SEC) using a Superose 6 Increase 3.2/300 (Cytiva) with buffer D. The major peak fraction was further concentrated (100 kD cut-off Eppendorf concentrator, Amicon) and directly used for SAXS analysis.

### SAXS

We collected the SAXS data from stalled Hly T1SS on beamline BM29 at the ESRF Grenoble (76). The BM29 beamline was equipped with PILATUS 3 × 2M detector (Dectris) at a fixed distance of 2.813 m. The measurements were performed at 10°C with a protein concentration of 0.50 mg/ml. We collected 10 frames with an exposure time of 1 sec/frame and scaled the data to absolute intensity against water. All used programs for data processing were part of the ATSAS Software package (Version 3.0.5) (77). Primary data reduction was done with the program PRIMUS (78). The Guinier approximation (79) was used to determine the forward scattering *I*(*0*) and the radius of gyration (*R*_g_). The pair-distribution function *p(r)* was created with the program GNOM (80) and determined the maximum particle dimension (*D_max_*). Ab initio models were created with DAMMIF (81) and the superpositioning was done with SUPCOMB (82).

### Data, materials, and software availability

All study data are included in the article and/or SI. For molecular simulations, the AMBER22 package of molecular simulation codes was used. AMBER22 is available from here: http://ambermd.org/.

## Acknowledgments

This study was funded by the Deutsche Forschungsgemeinschaft (DFG, German Research Foundation), Project 267205415/CRC1208, granted to L.S. (subproject A01) and H.G. (subproject A03) and, in part, by a grant from the Ministry of Innovation, Science, and Research within the framework of the NRW Strategieprojekt BioSC (No. 313/323-400-00213). The Center for Structural Studies is funded by the DFG (Grant number 417919780 and INST 208/761-1 FUGG to S.S.). We thank Filip König (HHU Düsseldorf) for the help with changes in the Constraint Network Analysis software to accommodate large molecular systems. We are grateful for the computational support by the “Zentrum für Informations und Medientechnologie” at the Heinrich Heine Universität Düsseldorf and the Gauss Centre for Supercomputing e.V. (www.gauss-centre.eu) for funding this project by providing computing time to H.G. through the John von Neumann Institute for Computing (NIC) on the GCS Supercomputer JUWELS (83) at Jülich Supercomputing Centre (JSC) (user ID: t1ss). We would like to thank all members of the Institute of Biochemistry for stimulating discussions and especially Manuel Anlauf for cloning the plasmid. We acknowledge the European Synchrotron Radiation Facility (ESRF) for the provision of synchrotron radiation facilities, and we would like to thank Petra Pernot for assistance in using beamline BM29. We also acknowledge DESY (Hamburg, Germany), a member of the Helmholtz Association HGF, for the provision of experimental facilities. Parts of this research were carried out at PETRA III, and we would like to thank Cy M. Jeffries and Dmytro Soloviov (EMBL Hamburg) for assistance in using beamline P12.

## Notes

### Competing Interest Statement

The authors have declared no competing interest.

