## Supporting Information for "Molecular Insights into CLD Domain Dynamics and Toxin Recruitment of the HlyA *E. coli* T1SS"

* Prof. Dr. Holger Gohlke

Institute for Pharmaceutical and Medicinal Chemistry, Heinrich Heine University Düsseldorf, Universitätstr. 1. 40225 Düsseldorf, Germany
and
Institute of Bio- and Geosciences (IBG-4: Bioinformatics), Forschungszentrum Jülich GmbH, Wilhelm-Johnen-Str., 52425 Jülich, Germany

**Supporting Text**

*ATP parametrization*

Partial atomic charges and parameters for ATP, compatible with the GAFF2 force field, were derived using a multi-stage, fragment-based approach using the AmberTools (1) software package. The ATP molecule was split into three capped building blocks: *N*9-Methyladenine, Methyl triphosphate, and 1-(β-D-Ribofuranosyl)imidazole. To capture conformational variability due to sugar puckering, two distinct conformers of the 1-(β-D-Ribofuranosyl)imidazole fragment were included.

Initial geometries for each fragment conformer were optimized using Gaussian 16 (2) at the Hartree-Fock (HF) level of theory with the 6-31G* basis set, applying tight convergence criteria. In order to adhere to best practices for generating a robust molecular electrostatic potential (MEP), a rigid body reorientation algorithm, inspired by protocols used by the R.E.D. Server (3), was used to generate two distinct spatial orientations for each optimized conformer.

The MEP was subsequently calculated for every structure (conformer and orientation) at the HF/6-31G* level using the Merz-Kollman scheme. Standard grid parameters were employed, consisting of four concentric layers constructed with van der Waals radii scaling factors of 1.4, 1.6, 1.8, and 2.0, and a surface point density of 1 point per $Å^2$. Atom types were assigned according to the GAFF2 specification using antechamber. A two-stage Restrained Electrostatic Potential (RESP) (4) fitting was performed, utilizing the ensemble of MEP data derived from all conformers and orientations for each fragment. During this process, charge constraints defined the fragment boundaries at the capping groups, and charge equivalencing was enforced for chemically symmetric atoms, following established fragmentation procedures (1). Bond, angle, or dihedral parameters absent from the standard GAFF2 parameter set were identified and generated using parmchk2.

**Supporting Figures**

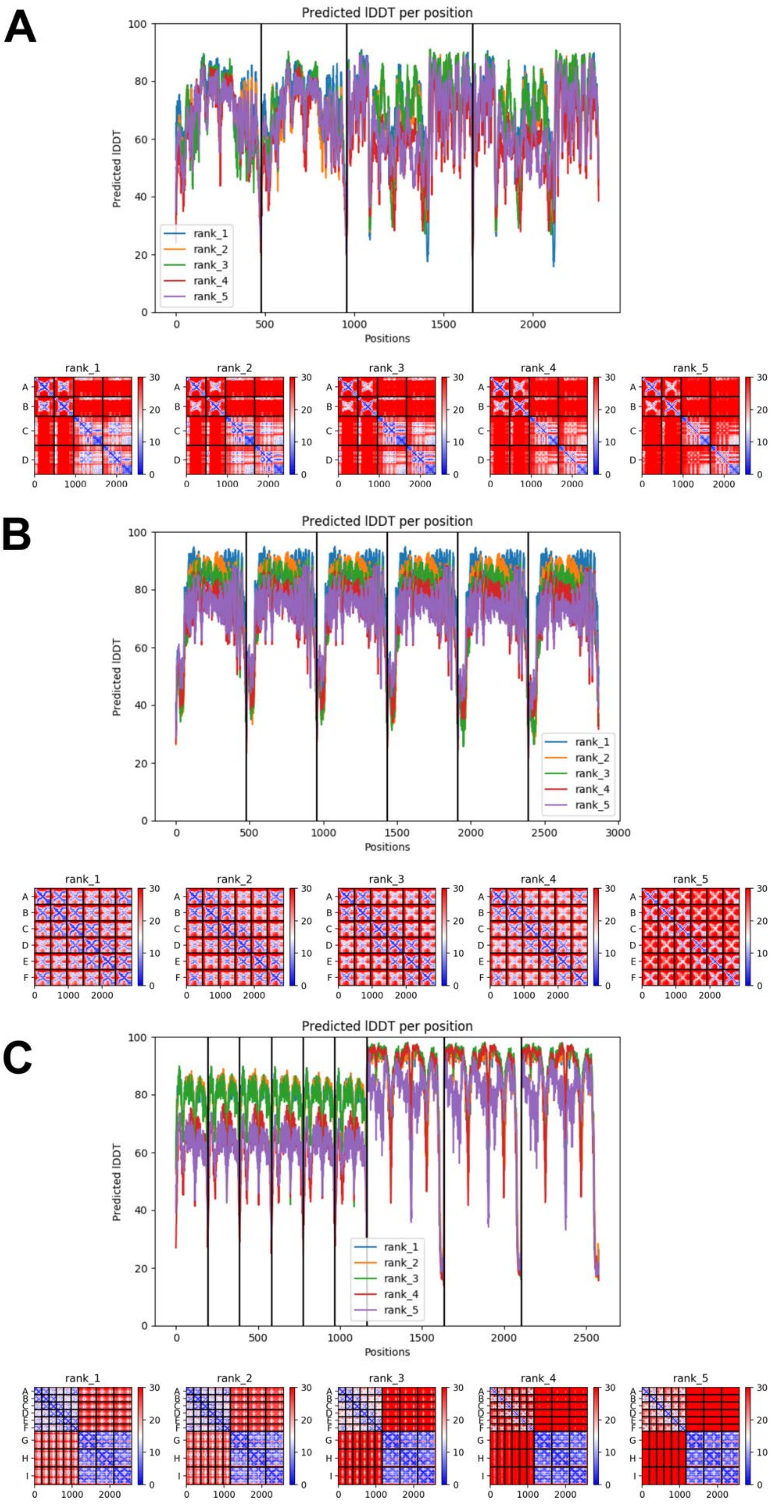


**Figure S1. pLDDT (top) and Predicted Alignment Error (PAE, bottom) scores for the generated models. A)** 2xHlyB$_O$/2xHlyD model; **B)** 6xHlyD and **C)** 6xHlyD (residues 132-385)/3xTolC. The model ranked the best was always selected for subsequent modeling steps.

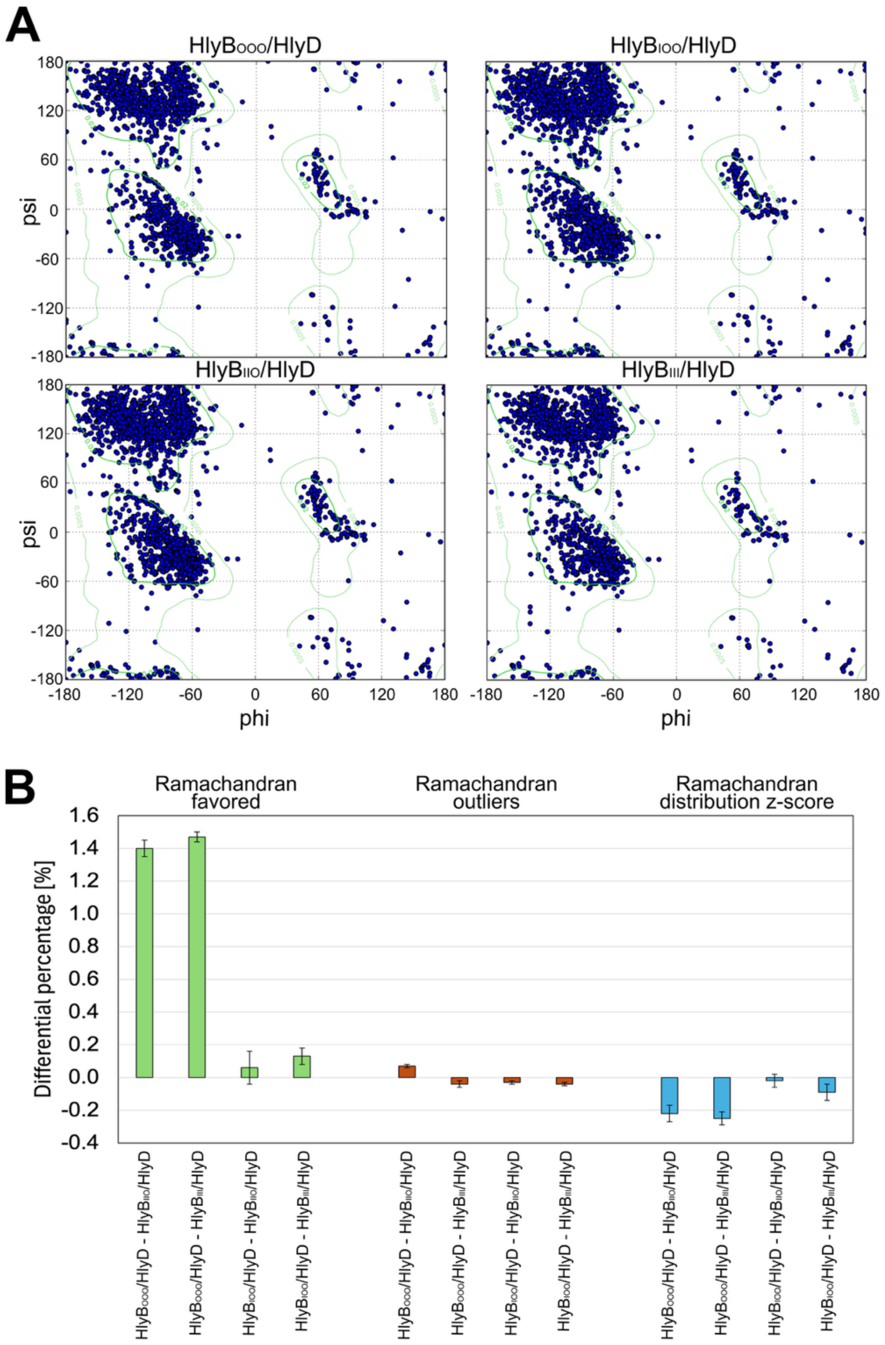


**Figure S2. Ramachandran plot analysis of HlyB/HlyD complexes. A)** Ramachandran plots computed with MolProbity for four modeled systems $HlyB_{OOO}$/HlyD, $HlyB_{IOO}$/HlyD, $HlyB_{IIO}$/HlyD and $HlyB_{III}$/HlyD. Blue dots indicate $\phi/\psi$ angles for individual residues; green contours represent favored regions. **B)** Comparative analysis of Ramachandran statistics across systems. Bar plots show differential percentages for residues in favored regions (green), outliers (orange) and Ramachandran distribution Z-scores (cyan), calculated relative to the $HlyB_{IIO}$/HlyD and $HlyB_{III}$/HlyD with respect to the $HlyB_{OOO}$/HlyD, $HlyB_{IOO}$/HlyD (configuration present in the cryo-EM structures). Error bars represent the standard error of each calculation.

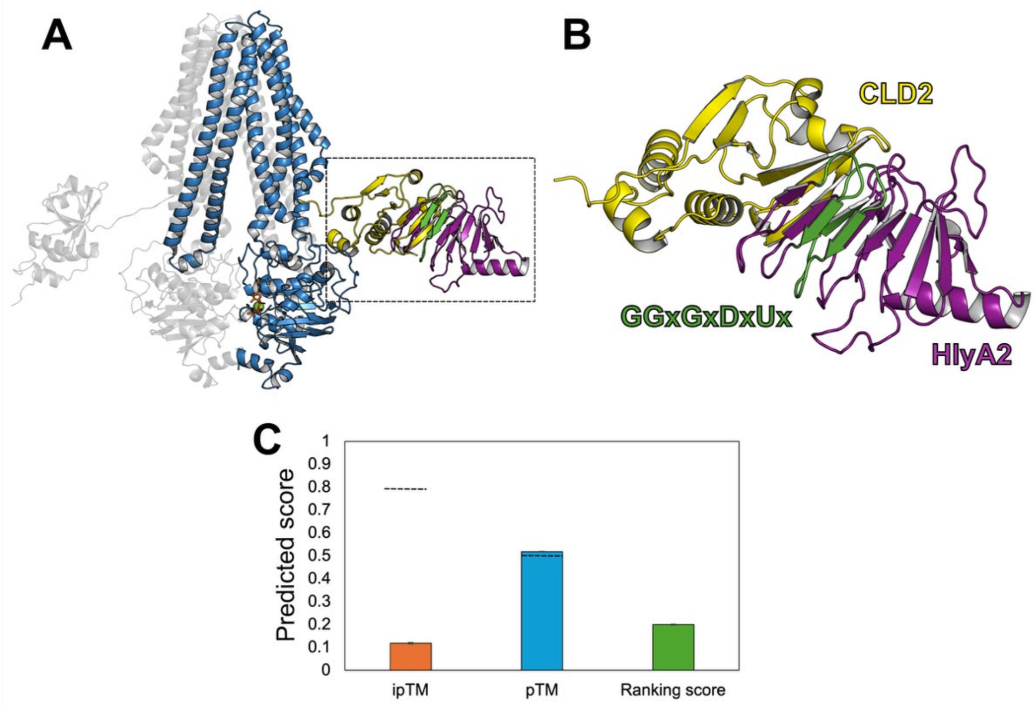


**Figure S3. Structural model of HlyA2 bound to CLD2 of HlyB predicted with AlphaFold3** (6). **A)** The HlyA2/HlyB$_{IOO}$ complex is depicted. Only one inward-facing HlyB monomer of HlyB$_{IOO}$ is shown as blue ribbons, and the CLD2 motif is shown as yellow ribbons. The predicted HlyA2 is shown as purple and green ribbons. **B)** Close-up view of the CLD2-HlyA2 complex. The GG-motifs known to interact with HlyA2 are modeled as β-sheets according to the predicted (6) secondary structure. **C)** Model scores indicating that the predicted complex quality is low. The interface predicted template modeling (ipTM) score describes the model quality of the complex interface. Values below 0.8 (dotted line) likely suggest a failed prediction (7). pTM is an integrated measure of how well the overall complex structure of the complex is predicted. A pTM score below 0.5 indicates the predicted structure is likely wrong (7). The ranking score is a composite score, used to sort and rank multiple predicted models. A ranking score ~0.9 or higher is typically considered for a high-confidence model (6).

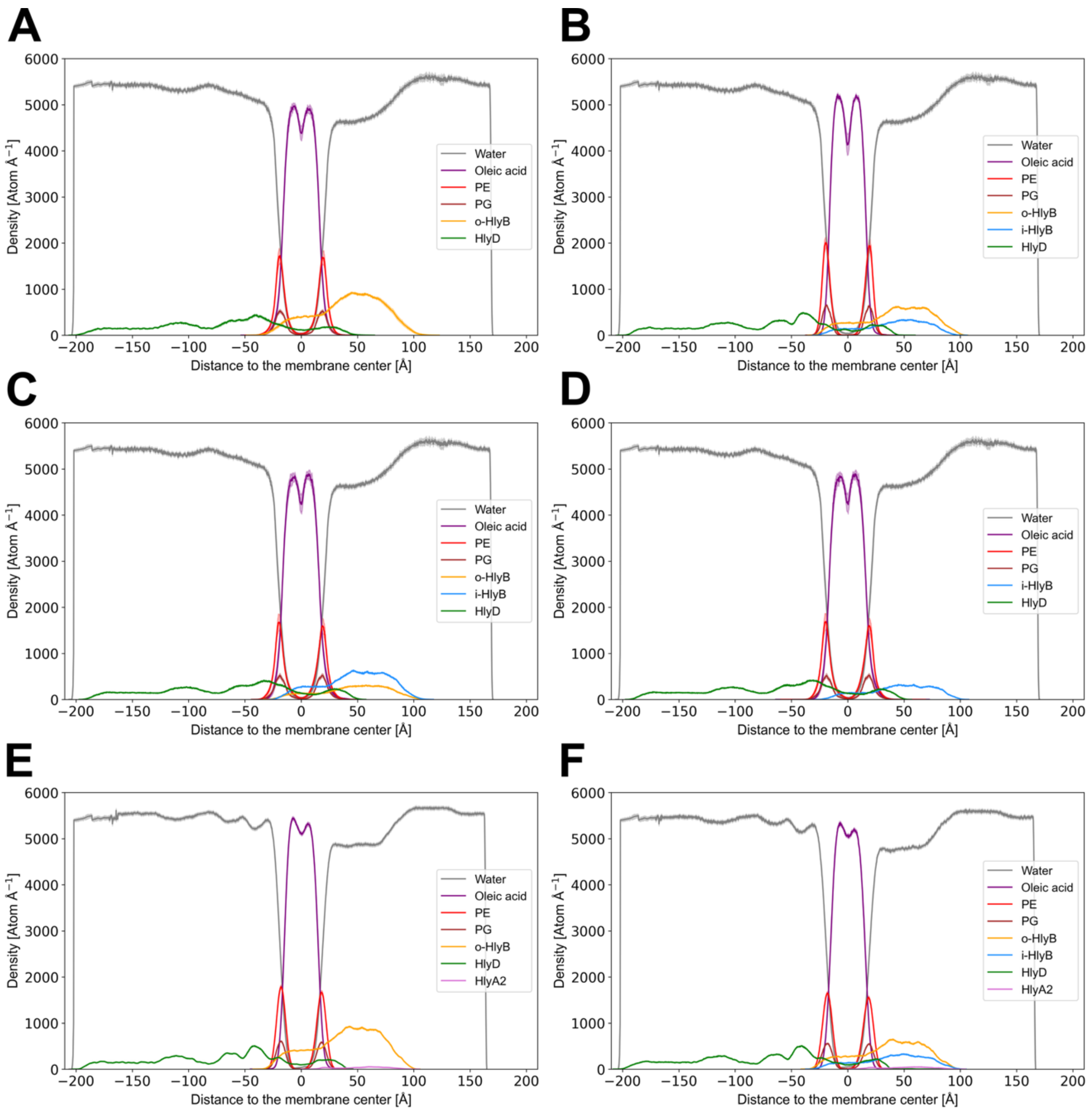


**Figure S5.** Atom density profiles of membrane components averaged over five independent unbiased MD simulations of **A)** HlyB$_{OOO}$/HlyD, **B)** HlyB$_{IOO}$/HlyD, **C)** HlyB$_{IIO}$/HlyD, **D)** HlyB$_{III}$/HlyD, **E)** HlyA2/HlyB$_{OOO}$/HlyD, and **F)** HlyA2/HlyB$_{IOO}$/HlyD (Table S1). The obtained shapes correspond with those generally found by experiments and MD simulations for membranes of Gram-negative bacteria (5).

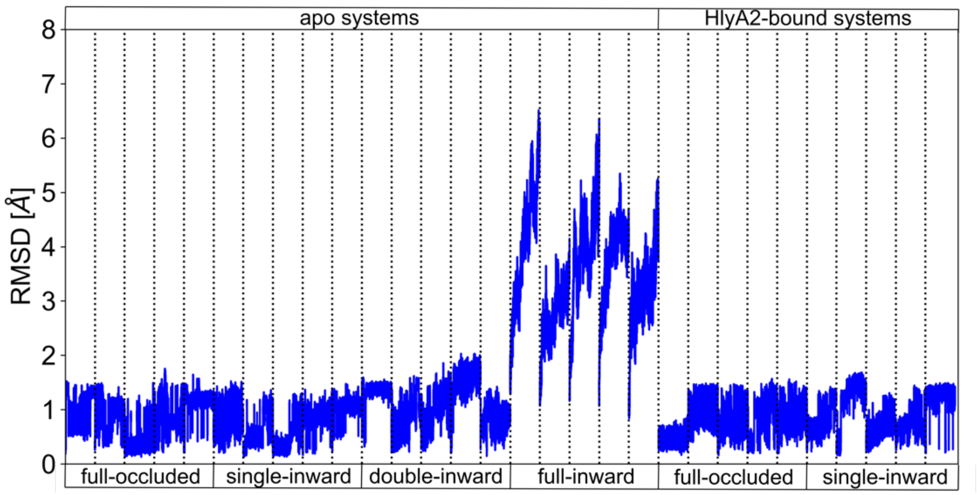


**Figure S6. Structural variability of adenosine triphosphate (ATP)-Mg$^{2+}$ moieties** in MD simulations of apo HlyB$_{OOO}$/HlyD, HlyB$_{IOO}$/HlyD, HlyB$_{IIO}$/HlyD, and HlyB$_{III}$/HlyD as well as HlyA2/HlyB$_{OOO}$/HlyD and HlyA2/HlyB$_{IOO}$/HlyD (Table S1). In all replicas besides the full-inward apo system, the ATP-Mg$^{2+}$ moieties preserve their position after 1 µs with respect to the first frame (RMSD < 2 Å). The RMSD of all ligand atoms was computed after superpositioning the C$_{\alpha}$ atoms of HlyB. Dotted lines separate independent replicas.

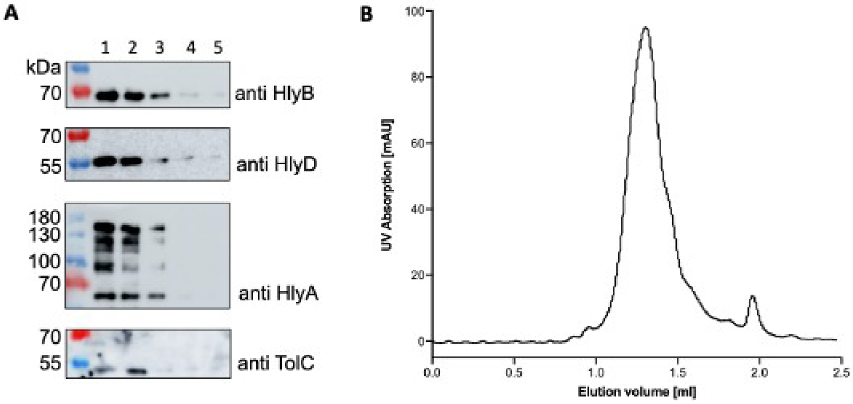


**Figure S7.** Purification of the entire HlyA T1SS. A) eGFP-stalled T1SS was purified via flag-tag resin. Western blot analysis confirming the presence of HlyB, HlyD, HlyA, and TolC in the elution fractions. B) Chromatogram representing the purification of the stalled HlyA T1SS by size-exclusion chromatography. The y-axis represents the UV absorption of the protein at 280 nm, while the x-axis represents the elution volume.

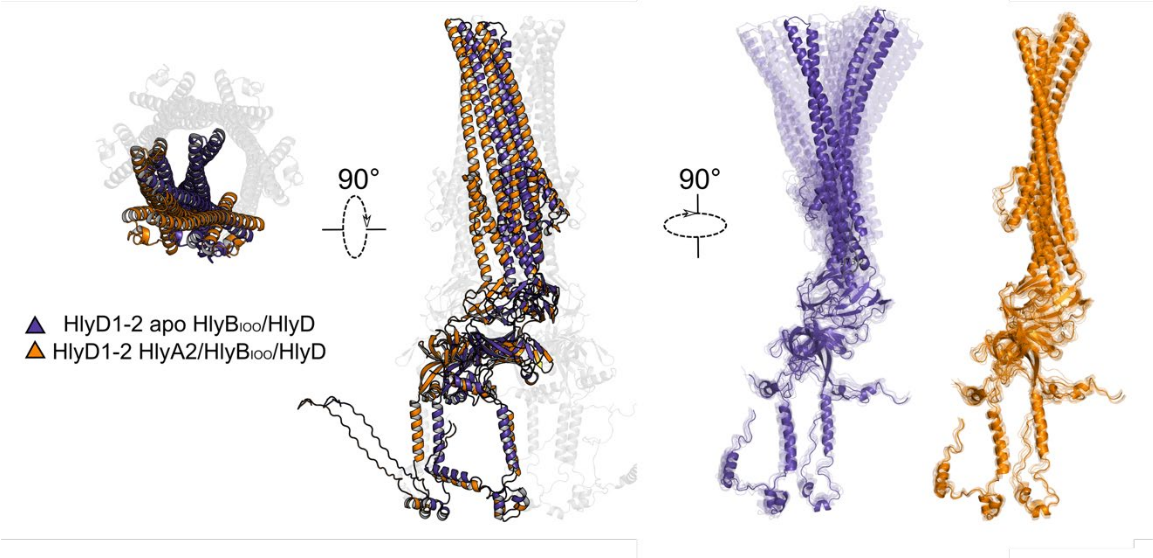


**Figure S8.** Structural overlay of configurations representing the average structures of HlyD1 and HlyD2 for the apo (purple ribbon) and HlyA2-bound (orange ribbon) HlyB$_{IOO}$/HlyD complexes. The overlay is shown from the extracellular side and depicted with z-clipping at the level of 150 Å from the top (left) and as a frontal view (center). The remaining HlyD chains are represented in grey ribbons. Structures obtained by MD simulations show higher HlyD1-2 mobility in the apo structures (purple) compared to the HlyA2-bound structures (orange).

**Supporting Tables**

**Table S1.** Systems generated with Packmol-Memgen and subjected to MD simulations and analyses.

| System | HlyB configuration[a] | Inner membrane composition | Number of inward-facing HlyB protomers | HlyA2 bound to | Replicas | Simulation length per replica [µs] |
|---|---|---|---|---|---|---|
| $HlyB_{OOO}$/HlyD | Full-occluded | DOPE:DOPG 3:1 | 0 | - | 5 | 1 |
| $HlyB_{IOO}$/HlyD | Single-inward | DOPE:DOPG 3:1 | 1 | - | 5 | 1 |
| $HlyB_{IIO}$/HlyD | Double-inward | DOPE:DOPG 3:1 | 2 | - | 5 | 1 |
| $HlyB_{III}$/HlyD | Full-inward | DOPE:DOPG 3:1 | 3 | - | 5 | 1 |
| HlyA2/CLD | - | - | - | CLD | 5 | 1 |
| HlyA2/$HlyB_{OOO}$/HlyD | Full-occluded | DOPE:DOPG 3:1 | 0 | CLD2 of $HlyB_{O}$ | 5 | 1 |
| HlyA2/$HlyB_{IOO}$/HlyD | Single-inward | DOPE:DOPG 3:1 | 1 | CLD2 of $HlyB_{I}$ | 5 | 1 |

[a] All simulated HlyB protomers are bound to ATP and $Mg^{2+}$ except for (V).

**Table S2.** Oligonucleotides used in this study.

| Oligonucleotide | 5' → 3' sequence |
|---|---|
| HlyA-L714-FLAGx3-L-fw | ACAAGGACGACGATGATAAATATTCCGTGGAAGAACTTATTG |
| HlyA-L714-FLAGx3-L-rev | TCGTCATCATCTTTATAATCTAAGTTATCAGTCTCTGTTAAATTTTTAC |
| FLAGx3-L (single-stranded oligo) | GATTATAAAGATGATGACGACAAGGGTGGCGGCGGTTCTGACTACAAGGACGAT GATGACAAGGGCGGTGGCGGTTCTGACTACAAGGACGACGATGATAAA |

**Table S3.** Overall SAXS data.

| | |
|---|---|
| **Data collection parameters** | |
| SAXS Device | BM29, ESRF Grenoble (8) |
| Detector | PILATUS 3 X 2M |
| Detector distance (m) | 2.813 |
| Beam size | 200 µm x 100 µm |
| Wavelength (nm) | 0.099 |
| Sample environment | Quartz glass capillary, 1 mm ø |
| Absolute scaling method | Comparison with scattering from pure $H_2O$ |
| Normalization | To transmitted intensity by beam-stop counter |
| Scattering intensity scale | Absolute scale, $cm^{-1}$ |
| *s* range ($nm^{-1}$)‡ | 0.06 – 5.0 |
| **Sample** | **HlyA stalled T1SS** |
| Organism | *E. coli UTI 89* |
| UniProt ID | Q1R2T6 (HlyB), Q1R2T7 (HlyD), Q1R2T5 (HlyA), P02930 (TolC) |
| Mode of measurement | batch |
| Temperature (°C) | 10 |
| Exposure time (# frames) | 1 (10 Frames) |
| Protein buffer | 50 mM Tris pH 7.5, 150 mM NaCl, 10 mM CaCl2, 0.0063% GDN |
| Protein concentration (mg/ml) | 0.50 |
| **Structural parameters** | |
| *Guinier Analysis (PRIMUS)* | |
| $I(0) \pm \sigma$ ($cm^{-1}$) | 1042.61 ± 7.68 |
| $R_g \pm \sigma$ (nm) | 11.35 ± 0.10 |
| *s-range* ($nm^{-1}$) | 0.063 – 0.114 |
| *min < sRg < max limit* | 0.713 – 1.298 |
| Data point range | 1 - 11 |
| Linear fit assessment ($R^2$) | 0.997 |
| *PDDF/P(r) Analysis (GNOM)* | |
| $I(0) \pm \sigma$ ($cm^{-1}$) | 1070.00 ± 6.75 |
| $R_g \pm \sigma$ (nm) | 12.17 ± 0.099 |
| $D_{max}$ (nm) | 46.20 |
| Porod volume ($nm^3$) | 3596.02 |
| *s-range* ($nm^{-1}$) | 0.0629 – 1.990 |
| χ2 / CorMap P-value | 0.978 / 0.998 |
| **Molecular mass (kDa)** | |
| From *I*(0) | 1042.61 |
| From MoW2 (9) | 1086.06 |
| From Size & Shape (10) | 1106.61 |
| Bayesian Inference (11) | 964.85 |
| From sequence | 1106 (1x HlyA:6x HlyB:6x HlyD:3x TolC) Without detergent |
| **Ab-initio modeling** | |
| DAMMIF | |
| Symmetry | P3 |
| *s*-range for fit ($nm^{-1}$) | 0.063 – 1.99 |
| $\chi^2$, CorMap *P*-value | 1.071 / 0.519 |
| **SASBDB accession codes** (12) | XXX [This will be added later after upload of the data.] |
| **Software** | |
| ATSAS Software Version (13) | 3.0.5 |
| Primary data reduction | PRIMUS (14) |
| Data processing | GNOM (15) |
| *Ab initio* modeling | DAMMIF (16) |
| Model visualization | PyMOL (17) |
